# Amantadine modulates action-specific neural ensembles in hypokinetic and hyperkinetic conditions

**DOI:** 10.1101/2025.04.01.646527

**Authors:** Gaurav Chattree, Radosław Chrapkiewicz, Yanping Zhang, Jane Li, Fatih Dinc, Mark J. Schnitzer

**Author notes:** Present address: Kavli Institute for Theoretical Physics, UC Santa Barbara, Santa Barbara CA 93106 USA. **Correspondence to:** Dr. Gaurav Chattree, Department of Neurology & Neurological Sciences, Stanford University, 213 Quarry Road, Palo Alto, CA 94304.; Dr. Mark J. Schnitzer, Departments of Biology, Applied Physics & Neurosurgery, HHMI, Stanford University, 318 Campus Drive, Stanford, CA 94305.

## Abstract

**Background:** In the classical model of basal ganglia circuitry, striatal spiny projection neurons of the direct and indirect pathways (dSPNs, iSPNs) promote and suppress movement, respectively, and exhibit unbalanced activity levels during hypokinetic or hyperkinetic conditions. Most therapies for these conditions are thought to work by rebalancing the relative activity of dSPNs and iSPNs towards normal levels. However, the mechanism of amantadine, which uniquely improves both hypokinetic and hyperkinetic conditions, is poorly understood.

**Objective:** To determine whether amantadine restores motor function by normalizing the balance of dSPN and iSPN activity or through a distinct mechanism.

**Methods:** We used dual-color two-photon Ca^2+^ imaging in the 6-OHDA mouse model of Parkinson’s disease to concurrently monitor dSPN and iSPN dynamics across healthy, hypokinetic (parkinsonian), and hyperkinetic (dyskinetic) conditions.

**Results:** We evaluated both dSPN/iSPN activity balance and action-specific neural ensemble activity in the dorsolateral striatum. In hypokinetic conditions, L-DOPA rescued the dSPN/iSPN imbalance but failed to restore the disrupted activity of the locomotion-specific ensemble. Conversely, amantadine improved locomotion-specific ensemble activity without normalizing the dSPN/iSPN imbalance. In hyperkinetic conditions, forelimb dyskinesias were characterized by neural activity patterns distinct from those encoding locomotion. Amantadine selectively suppressed the resting activity of forelimb dyskinesia ensembles without affecting locomotion-coding ensembles or restoring pathway balance.

**Conclusions:** L-DOPA and amantadine may act through distinct mechanisms, with L-DOPA normalizing pathway balance and amantadine modulating action-specific neural ensembles. These findings support the importance of action-coding disruptions during hypokinetic and hyperkinetic conditions and suggest that correcting them can restore motor function.

## INTRODUCTION

In the classical model of basal ganglia function, the two major anatomical pathways of the basal ganglia have opposing roles in the regulation of movement. According to the model, the activation of striatal spiny projection neurons of the direct pathway (dSPNs) elicits movement, whereas the activation of spiny projection neurons of the indirect pathway (iSPNs) suppresses movement.^1–3^ These classical conceptions provide a basic framework for understanding movement disorders, which are broadly classified into hypokinetic disorders characterized by reduced movement, such as Parkinson’s disease, and hyperkinetic disorders characterized by excessive involuntary movement, such as L-DOPA-induced dyskinesias and Huntington’s disease.^1,2,4–7^ Physiological recordings and SPN-type-specific manipulations in animal models of these conditions support this model, showing that increased dSPN activity and/or decreased iSPN activity contributes to hyperkinetic states, whereas decreased dSPN activity and/or increased iSPN activity contributes to hypokinetic states.^1,4,8–14^ However, this model does not fully capture the complexity of how the striatum contributes to movement control.^15^ In normal brain states, dSPNs and iSPNs exhibit approximately equal activity levels across the initiation, performance, and offset of movement, and this coordination may be important for well-choreographed motor control.^16–18^ The striatum also exhibits action-specific patterns of neural activation, in which distinct subsets of dSPNs and iSPNs jointly form ensembles to encode particular actions.^8,19^ This suggests that action-specific ensemble coding is an additional, distinct aspect of striatal physiology.

Most treatments for movement disorders can be understood in terms of modulating pathway balance. Hyperkinetic disorders are treated using drugs intended to decrease dSPN activity and/or increase iSPN activity, while hypokinetic disorders are treated conversely.^4,11,20^ However, amantadine, a clinically unique medication that improves both hypokinetic and hyperkinetic conditions, has a poorly understood mechanism of action.^21–24^ Notably, it is unclear whether it works by restoring pathway balance or by acting on other aspects of striatal physiology.

Resolving this question could guide therapeutic development. While clinically useful, amantadine has variable efficacy and dose-limiting side effects.^21^ A therapy with a similar, but more precise, mechanism could be highly valuable for treating a broad number of movement disorders. However, development of such a therapeutic has been limited by amantadine’s broad pharmacology and lack of selectivity across numerous receptor targets.^21–28^ Given amantadine’s broad activity profile it may have complex actions on neural circuits.

To probe amantadine’s circuit-level effects, we imaged dSPN and iSPN dynamics concurrently in behaving mice across normal, hypokinetic, and hyperkinetic conditions before and after treatments comprising L-DOPA, amantadine, or a combination of both drugs. We evaluated the hypokinetic condition by using the 6-OHDA model of Parkinson’s disease and the hyperkinetic condition via L-DOPA-induced dyskinesias.^29^ In these conditions, we evaluated both the relative balance of dSPN/iSPN activity and the activity of action-specific ensembles. We found that L-DOPA and amantadine act separably on these two aspects of striatal physiology. In the hypokinetic state, L-DOPA rescued the dSPN/iSPN imbalance but failed to restore the disrupted activity of the locomotion-specific ensemble. Conversely, amantadine improved locomotion ensemble coding without correcting the dSPN/iSPN imbalance. In the hyperkinetic state, amantadine selectively suppressed the resting activity of the dyskinesia ensembles while sparing the locomotion ensembles, again without restoring pathway balance.

Together, these results suggest two complementary therapeutic mechanisms: pathway rebalancing by L-DOPA, and neural ensemble modulation by amantadine. This framework provides insight into amantadine’s unique clinical profile and may guide future movement-disorder therapeutic development.

## METHODS

### Neural Ca^2+^ Imaging in the Dorsolateral Striatum of Awake, Head-Fixed Mice

Details regarding all experimental procedures and data analysis are provided in the **Supplemental Methods.** To examine the activity of dSPNs and iSPNs concurrently, we used dual-color two-photon microscopy to image somatic SPN Ca^2+^ dynamics in head-fixed behaving mice. We used Drd1a-Cre transgenic mice, crossed with Ai14 Cre-dependent mice, to selectively label dSPNs with the red fluorophore tdTomato.^30,31^ We then injected AAV2/9-CaMKII-jGCaMP7f virus into the dorsolateral striatum (DLS), driving expression of the green fluorescent Ca^2+^ indicator jGCaMP7f in both dSPNs and iSPNs.^32^ We focused on the DLS, as it is implicated in the pathophysiology of parkinsonism and L-DOPA-induced dyskinesia.^9,13,33,34^ We used the CaMKII promoter, as it labels both SPN populations while minimizing expression in striatal interneurons.^8,35,36^

To gain optical access to the striatum for two-photon imaging, about a week after virus injection we implanted a gradient index (GRIN) lens over the DLS.^8,37–39^ In each mouse, we recorded Ca²⁺ activity at 3 different tissue planes during each imaging session. We imaged each plane for a total duration of 20.0 min per session, parsed into two non-successive blocks of 10.0 min each, yielding a total of 60.0 min of Ca^2+^-imaging data per session. Since dSPNs co-expressed red tdTomato and green jGCaMP7f, whereas iSPNs expressed only jGCaMP7f, dual-color two-photon imaging allowed us to distinguish these two cell types (**Figures 1A,B**). On average, we identified 371 ± 119 (mean ± s.d.) tdTomato-positive dSPNs per session, and 285 ± 179 GCaMP-positive, tdTomato-negative iSPNs per session, which led to a mean dSPN-to-iSPN cell count ratio of 1.7 ± 1.0 per session (n = 63 sessions from 6 mice). However, despite more dSPNs being identified overall, most cells with detectable Ca²⁺ activity were iSPNs (78 ± 11%). We suspect this imbalance may reflect the possibility that the use of the Drd1a-Cre driver mouse line to express tdTomato in dSPNs hindered the expression of GCaMP in these cells, as a similar imbalance has been observed before with this strategy.^8^ To reduce the extent to which this unequal cell count biased our pathway-specific comparisons, we employed a 1:1 subsampling bootstrap procedure for all analyses comparing the relative activity of dSPNs and iSPNs (**Supp. Methods**), although we recognize that this does not fully resolve sampling bias in the detectable dSPN population.

**Figure 1.**
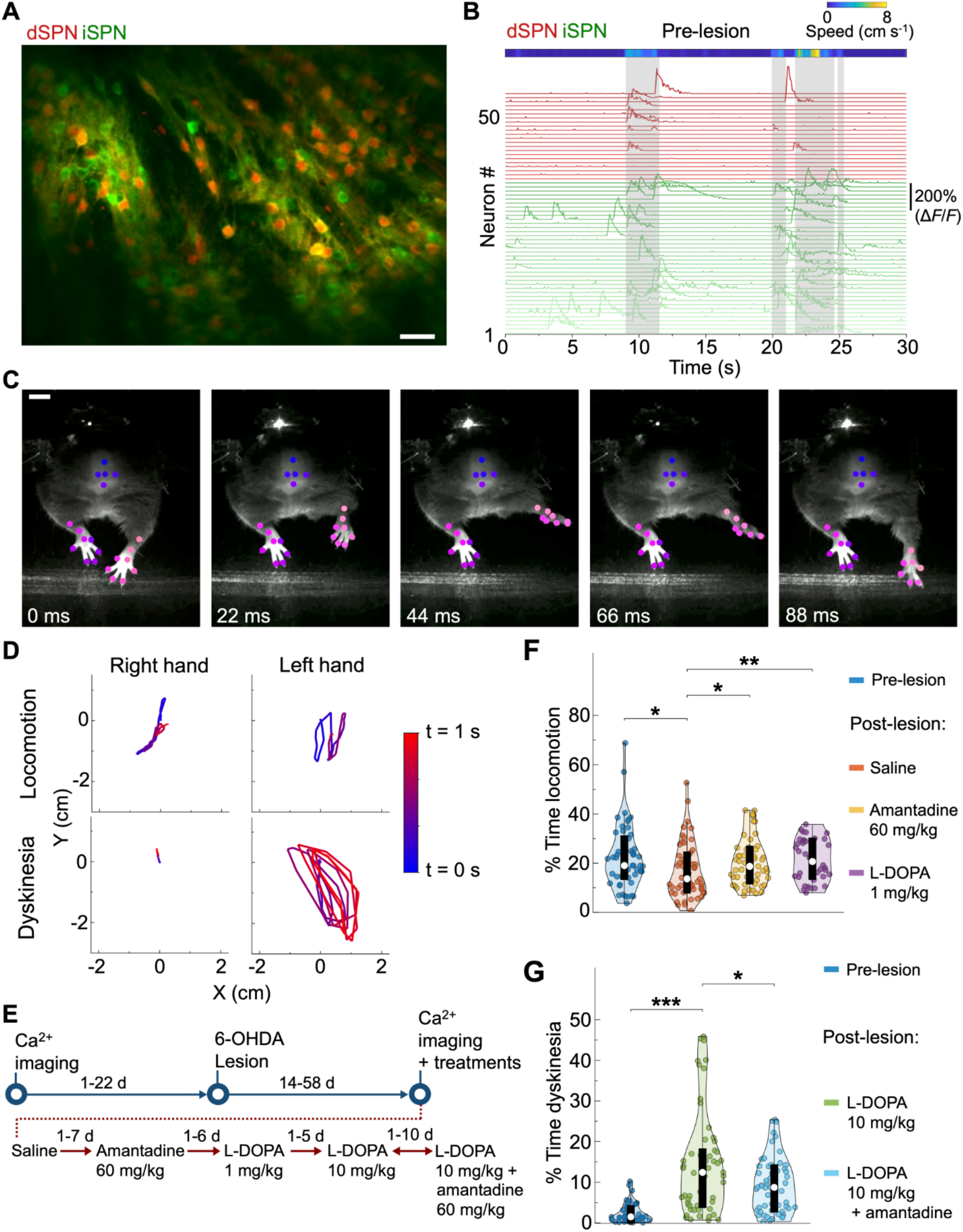
Imaging SPN Ca^2+^ activity and analysis of movement kinematics in awake mice across normal, hypokinetic, hyperkinetic, and treated conditions. (**A**) Example dual-color image of SPNs acquired by two-photon microscopy via a GRIN lens. dSPNs but not iSPNs expressed red fluorescent tdTomato. Both SPN-types expressed green jGCaMP7f. The image is a time-averaged projection of a movie lasting 10 min. Scale bar: 40 µm. **(B)** *Bottom*, Example traces of fluorescence Ca²⁺ activity (Δ*F*/*F*) of individual dSPNs (red traces) and iSPNs (green traces), recorded concurrently in a normal mouse. Gray shading marks periods of locomotion. *Top*, Color plot of the mouse’s locomotor speed on the wheel. **(C)** Representative sequence of image frames acquired by infrared videography, showing one phase of a rapid raising movement of the left forepaw, which was subsequently repeated (see **D**). This movement pattern was characteristic of L-DOPA-induced dyskinesia contralateral to a 6-OHDA lesion in the substantia nigra pars compacta (SNc). Colored dots denote body parts that were tracked using the machine vision analysis package, DeepLabCut.^40^ Different color dots denote different body parts (*e.g.*, right hand, left hand). Scale bar: 1 cm. **(D)** Traces of right and left forepaw movements (1-s-durations) in the two dimensions captured by infrared videography (90 fps) during representative bouts of locomotion (*top row*) and dyskinesia (*bottom row*). Color scale shows the time elapsed following movement initiation. **(E)** Experimental timeline for pre-lesion Ca^2+^ imaging and behavioral tracking sessions, 6-OHDA injection, and post-lesion Ca^2+^ imaging and behavioral testing sessions. We administered drug treatments in the order shown. After an imaging session with L-DOPA (1 mg/kg), sessions alternated between administration of either L-DOPA (10 mg/kg) or L-DOPA (10 mg/kg) + amantadine. **(F)** Violin plots showing the percentage of time in each block that mice spent in locomotion. Compared to before the 6-OHDA lesion, in the untreated hypokinetic condition we observed a decline in the time spent in locomotion, which was improved by both amantadine and L-DOPA (1 mg/kg) treatment. (*p < 0.05, **p < 0.01; one-sided rank sum tests, corrected for multiple comparisons using a Benjamini–Hochberg procedure with a false-discovery rate of 0.05; n = 50 pre-lesion blocks, 53 saline blocks, 53 amantadine blocks, and 38 L-DOPA (1 mg/kg) blocks from 5 mice). **Note:** Hypokinetic condition analyses use 5 mice due to exclusion of one animal with inadvertent L-DOPA exposure prior to its saline (control) session; hyperkinetic condition analyses (*e.g.*, **G**) use 6 mice (see **Methods**). **(G)** Violin plots of the percentage of time in each block that mice exhibited dyskinetic movement. The addition of amantadine to high-dose L-DOPA (10 mg/kg) led to reduced dyskinesia. The low levels of machine-vision-detected dyskinesias in pre-lesion blocks in actuality reflect grooming or other spontaneous non-locomotor movements. (*p < 0.05, ***p < 0.001; one-sided rank sum tests, corrected for multiple comparisons using a Benjamini–Hochberg procedure with a false-discovery rate of 0.05; n = 56 pre-lesion blocks, 57 L-DOPA (10 mg/kg) blocks, and 56 L-DOPA (10 mg/kg) + amantadine blocks from 6 mice). Here, and in violin plots throughout the paper, each violin shows a distribution of data for a given experimental condition, with the violin width denoting the probability density. The white dot in each violin marks the median value, the thick black box marks the interquartile range (IQR: 25th–75th percentiles), and the thin black lines extend to 1.5 times the IQR or the furthest data point within the IQR. Colored dots are individual data points.

### Behavioral Monitoring and Classification

To assess mouse movement during the Ca^2+^ imaging sessions, we head-fixed mice and allowed them to run on a wheel placed under the objective lens of the two-photon microscope. A rotary encoder reported rotations of the wheel, which we used to determine the mouse’s locomotor speed. An infrared camera captured the mouse’s forelimb movements. Machine vision analyses of the videos allowed us to identify forelimb dyskinesias, which we defined as rapid, choreiform forelimb movements that occurred independently of wheel motion (**Figures 1C,D, Supplemental Video 1**).^40^ We recorded neural activity and mouse behavior in each of the six different 10-min-blocks in each session. This allowed us to track activity in the two SPN populations concurrently across different brain states and treatment conditions **(Figure 1E)**.

### Analysis of Neural Activity Across Conditions

To enable rigorous comparisons across behavioral states, we focused our neural activity analyses on periods when mice were at rest. This approach offered two advantages. First, it reduced confounding behavioral variability across conditions in neural analyses. Second, in the seconds preceding movement, SPN activity patterns were predictive of upcoming actions. This allowed us to examine how the occurrence of these neural patterns at rest related to an animal’s overall propensity to move and how individual cells contributed to the encoding of upcoming actions.

## RESULTS

### Behavioral Phenotypes in Hypokinetic, Hyperkinetic, and Treated Conditions

To model a hypokinetic parkinsonian state, we unilaterally lesioned the substantia nigra pars compacta (SNc) with 6-hydroxydopamine (6-OHDA) (**Figure S1A**).^29^ The lesion decreased the proportion of time per block that mice engaged in locomotion as compared to pre-lesion baseline values. Treatment with either amantadine (60 mg/kg) or a standard dose of L-DOPA (1 mg/kg) led to improvements in the percentage of time spent locomoting per block, as compared to hypokinetic mice that received only saline (**Figure 1F**, **S2A**).

To model hyperkinetic conditions, we administered a high dosage of L-DOPA (10 mg/kg) to the same 6-OHDA-lesioned mice.^8^ This induced robust choreiform movements of the forelimb contralateral to the 6-OHDA lesion, consistent with L-DOPA-induced forelimb dyskinesias (**Figures 1C,D, Supplemental Video 1**). Administration of amantadine at 60 mg/kg, found previously to be effective at treating L-DOPA-induced dyskinesia in rodents,^25^ significantly reduced the percentage of time that mice exhibited forelimb dyskinetic movements when given together with high-dose L-DOPA (**Figure 1G**, **S2B**).

### L-DOPA and Amantadine Differentially Rescue Two Distinct Striatal Deficits in the Hypokinetic Condition

To identify the neural correlates of the hypokinetic condition and the mechanisms of therapeutic rescue, we investigated how L-DOPA and amantadine alter striatal dynamics. We first examined how SPN activity relates to movement across conditions by plotting the Ca²⁺ event rates of dSPNs and iSPNs relative to the times of locomotion onset and offset (**Figures S3A,B**).

Then, we evaluated the relative balance between the direct and indirect pathways by computing a dSPN-to-iSPN mean activity index (**Supp. Methods**). As expected, we found that the untreated hypokinetic (saline) condition was characterized by a significant reduction in this index (reflecting a relative shift toward indirect pathway activity) during both resting and pre-locomotion periods as compared to the pre-lesion baseline condition. The pre-locomotion period was defined as those 2-s-intervals without mouse movement, immediately prior to locomotion onset. Consistent with prior studies, treatment with L-DOPA (1 mg/kg) partially restored this pathway balance (**Figures 2A,B, S4A,B**).^8,11,13^ However, although amantadine also increased the percentage of time mice spent locomoting per block (**Figure 1F**), it failed to improve the dSPN-to-iSPN index (**Figures 2A,B)**. Consistent with this dissociation, both amantadine and L-DOPA (1 mg/kg) increased the probability of transitioning from rest to locomotion, irrespective of the dSPN-to-iSPN index value, and the dSPN-to-iSPN index showed no meaningful correlation with the speed or duration of locomotion bouts (**Figures S3C, S5A**). This discrepancy suggested that amantadine may act via a distinct mechanism independent of pathway balance. Therefore, we next examined another fundamental feature of striatal physiology, the activity of action-specific SPN ensembles.

**Figure 2.**
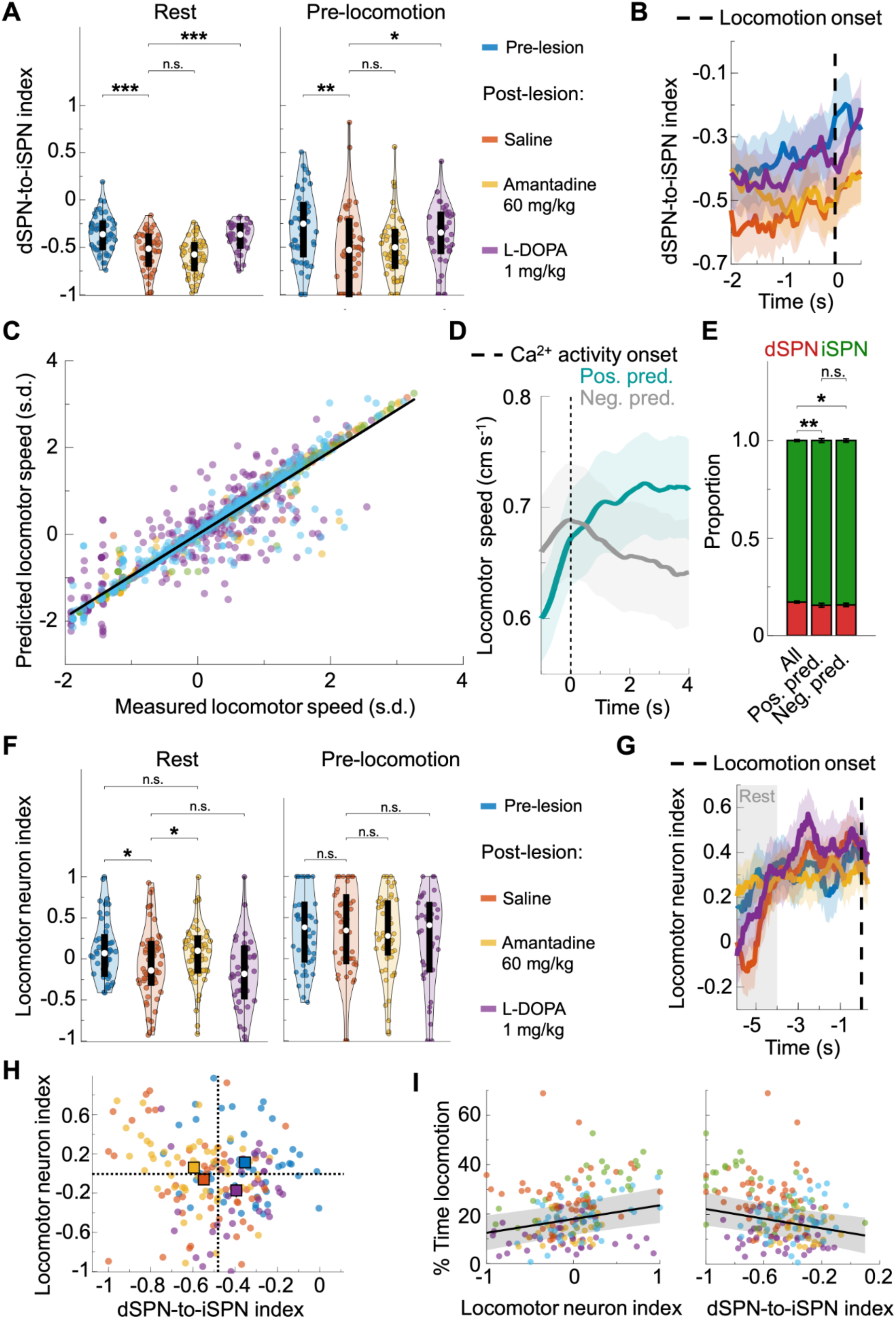
L-DOPA acts on dSPN to iSPN activity balance while amantadine acts on locomotion-specific ensembles during hypokinetic conditions. (**A**) Violin plots showing the distributions of dSPN-to-iSPN index values before and after the lesion, during resting (*left*) and pre-locomotion periods (*right*). L-DOPA (1 mg/kg) but not amantadine elevated the index values following the lesion. (*p < 0.05, **p < 0.01, ***p < 0.001; n.s. = not significant; two-sided rank sum tests corrected for multiple comparisons using a Benjamini–Hochberg procedure with a false-discovery rate of 0.05; n = 50 pre-lesion blocks, 53 saline blocks, 53 amantadine blocks, and 38 L-DOPA blocks from 5 mice). Color legend also applies to **B**. **(B)** Mean dSPN-to-iSPN index values plotted relative to locomotor onset time (vertical dashed line) for each condition. Colored shading: s.e.m. for n = 50 pre-lesion blocks, 53 saline blocks, 53 amantadine blocks, and 38 L-DOPA blocks from 5 mice. **(C)** Results of multivariate ridge regression analyses (for each mouse, imaging plane, and condition combination) predicting the speed of upcoming locomotion bouts based on the neural activity patterns just prior to movement. The scatter plot shows predicted *vs*. measured locomotion speeds for 5 mice, color-coded by mouse. Each data point shows results for an individual bout of locomotion. Speeds are z-scored for each mouse. Data is from pre-lesion, saline, amantadine, and L-DOPA (1 mg/kg) conditions and pooled here for visualization purposes. The overall regression line (black) has a training-set R² of 0.91 and a slope of 0.95. **(D)** Mean locomotion speed relative to the time of Ca²⁺ events for positive-predictive and negative-predictive locomotion neurons identified from the ridge regression model of (**C**), averaged across all Ca²⁺ events. Activation of positive-predictive (teal) or negative-predictive (gray) locomotion neurons was associated with an increase or decrease in locomotion speed, respectively. Shading: s.e.m. for n = 193 experimental blocks comprising 50 pre-lesion blocks, 53 saline blocks, 53 amantadine blocks, and 37 L-DOPA (1 mg/kg) blocks from 5 mice (one L-DOPA block excluded from here and in any subsequent analyses requiring locomotion-predictive neurons because none were identified in that block). Dashed vertical line: Ca²⁺ event onset (*t* = 0). **(E)** Proportions of dSPNs and iSPNs across the sets of all neurons, positive-predictive locomotion neurons, and negative-predictive locomotion neurons (*p < 0.05, **p < 0.01; n.s. = not significant; two-sided rank sum tests corrected for multiple comparisons using a Benjamini–Hochberg procedure with a false-discovery rate of 0.05; n = 193 experimental blocks comprising 50 pre-lesion blocks, 53 saline blocks, 53 amantadine blocks, and 37 L-DOPA (1 mg/kg) blocks from 5 mice). Error bars: s.e.m. **(F)** Violin plots of the mean resting and pre-locomotion period locomotor neuron index per block, for 4 different brain states or treatment conditions. Amantadine but not L-DOPA (1 mg/kg) improved the resting locomotor neuron index. Statistical comparisons were performed using rank sum tests (*p < 0.05; n.s. = not significant; n = 50 pre-lesion blocks, 53 saline blocks, 53 amantadine blocks, and 37 L-DOPA (1 mg/kg) blocks from 5 mice). For the two significant comparisons, one-sided rank sum tests were used. For the non-significant (n.s.) comparisons, one-sided tests in each direction confirmed non-significance. Tests were corrected for multiple comparisons using a Benjamini–Hochberg procedure with a false-discovery rate of 0.05. **(G)** Mean locomotor neuron index values plotted relative to locomotor onset time (vertical dashed line) for each condition. Gray shading indicates a time period classified as rest. Colored shading: s.e.m. for n = 50 pre-lesion blocks, 53 saline blocks, 53 amantadine blocks, and 37 L-DOPA blocks from 5 mice. **(H)** Scatter plot of the mean resting period dSPN-to-iSPN index value (X-axis) versus the locomotor neuron index value (Y-axis). Each point represents a single experimental block, colored by condition (see legends in **A, F**). Large squares with error bars denote the mean ± s.e.m. for each condition. Relative to the baseline pre-lesion condition (blue, top-right), the post-lesion saline condition (orange, bottom-left) shows a deficit in both metrics. Note that L-DOPA treatment (purple; bottom-right) primarily restores the dSPN-to-iSPN index, whereas amantadine treatment (yellow; top-left) primarily improves the locomotor neuron index. n = 50 pre-lesion blocks, 53 saline blocks, 53 amantadine blocks, and 37 L-DOPA blocks from 5 mice. **(I)** Linear mixed-effects models show that the locomotor neuron index predicts the percentage of time spent locomoting (slope = 5.6, p < 0.001). By contrast, the dSPN-to-iSPN index was not a positive predictor of locomotion time. Points in the scatter plots denote data from individual blocks, color-coded by individual mice, with regression lines fit for the untreated hypokinetic (saline control) condition (n = 193 blocks from 5 mice).

To do this, we identified ensembles of neurons whose activity in the 4-s window prior to movement was predictive of subsequent locomotion speed using a multivariate ridge regression model, which provided a good fit to the training data (training-set R^2^ = 0.91) (**Figures 2C, S5B**). Shifting the 4-s-long predictor window further back in time (*e.g.*, to the interval [–5 s, –1 s] prior to locomotion) significantly reduced the models’ predictive performances (**Figure S5C**), suggesting that the [–4 s, 0 s] interval captured SPN activity most relevant to the upcoming locomotion. The model identified neurons with strongly positive or negative regression coefficients (z-score > 1 s.d.) which we classified as positive-predictive and negative-predictive locomotion neurons, respectively. We functionally validated these classifications: events from positive-predictive neurons were associated with a subsequent speed increase (one-sided signed-rank test, p < 0.001, n = 193 blocks from 5 mice), while events from negative-predictive neurons were associated with a decrease (one-sided signed-rank test, p = 0.03, n = 193 blocks from 5 mice) (**Supp. Methods**) (**Figure 2D**). These predictive ensembles were a mix of dSPNs and iSPNs, with both groups showing a similar composition to the overall recorded population (**Figure 2E**).

We then calculated a locomotor neuron index to quantify the balance between these positive-predictive and negative-predictive locomotion ensembles (**Supp. Methods**). This analysis revealed a second deficit: the locomotor neuron index was significantly reduced in the untreated hypokinetic state as compared to the baseline pre-lesion state (**Figures 2F,G**). Amantadine, but not L-DOPA, restored the locomotor neuron index to nearly pre-lesion levels during rest periods without a detectable effect during the immediate pre-locomotion period or the first second of locomotion behavior (**Figures 2F,G, S6A**, **S7A**).

Plotting both metrics together highlighted their dissociation: L-DOPA primarily shifted the dSPN-to-iSPN index, whereas amantadine primarily shifted the locomotor neuron index (**Figure 2H**).

Finally, we further assessed the behavioral relevance of the locomotor-coding ensembles using linear mixed-effects models. We found that the locomotor neuron index was a positive predictor of the time spent locomoting per block, as was the resting activity of the positive-predictive locomotion neurons themselves (**Figures 2I, S7B**). By contrast, the dSPN-to-iSPN index was not a positive predictor of locomotion (**Figure 2I**).

Together, these findings suggest that the hypokinetic state involves two distinct deficits: a pathway imbalance addressed by L-DOPA, and a suppression of locomotion-coding activity addressed by amantadine.

### Amantadine Selectively Suppresses the Dyskinesia-Predictive Ensemble Without Restoring Pathway Balance

We next investigated amantadine’s effects on neural activity in the hyperkinetic state. We found that the dyskinetic state induced by high-dose L-DOPA (10 mg/kg) was characterized by a significant increase in the resting dSPN-to-iSPN index, consistent with a hyperactive direct pathway as shown in previous studies (**Figures 3A,B, Figure S8A**).^8,11,13^ Co-administration of amantadine, which significantly reduced forelimb dyskinetic behavior (**Figure 1G**), did not improve this dSPN-to-iSPN imbalance, and instead elevated it further (**Figure 3B,C**). While resting periods with high dSPN-to-iSPN index values had modestly higher probabilities of transitioning from rest to dyskinesia than those with low dSPN-to-iSPN index values, the value of this index in the pre-dyskinesia period showed no meaningful correlation with upcoming dyskinesia speed or duration (**Figures S8B, S9A**). These findings suggest that amantadine’s anti-dyskinetic properties may not be mediated by a restoration of pathway balance.

**Figure 3.**
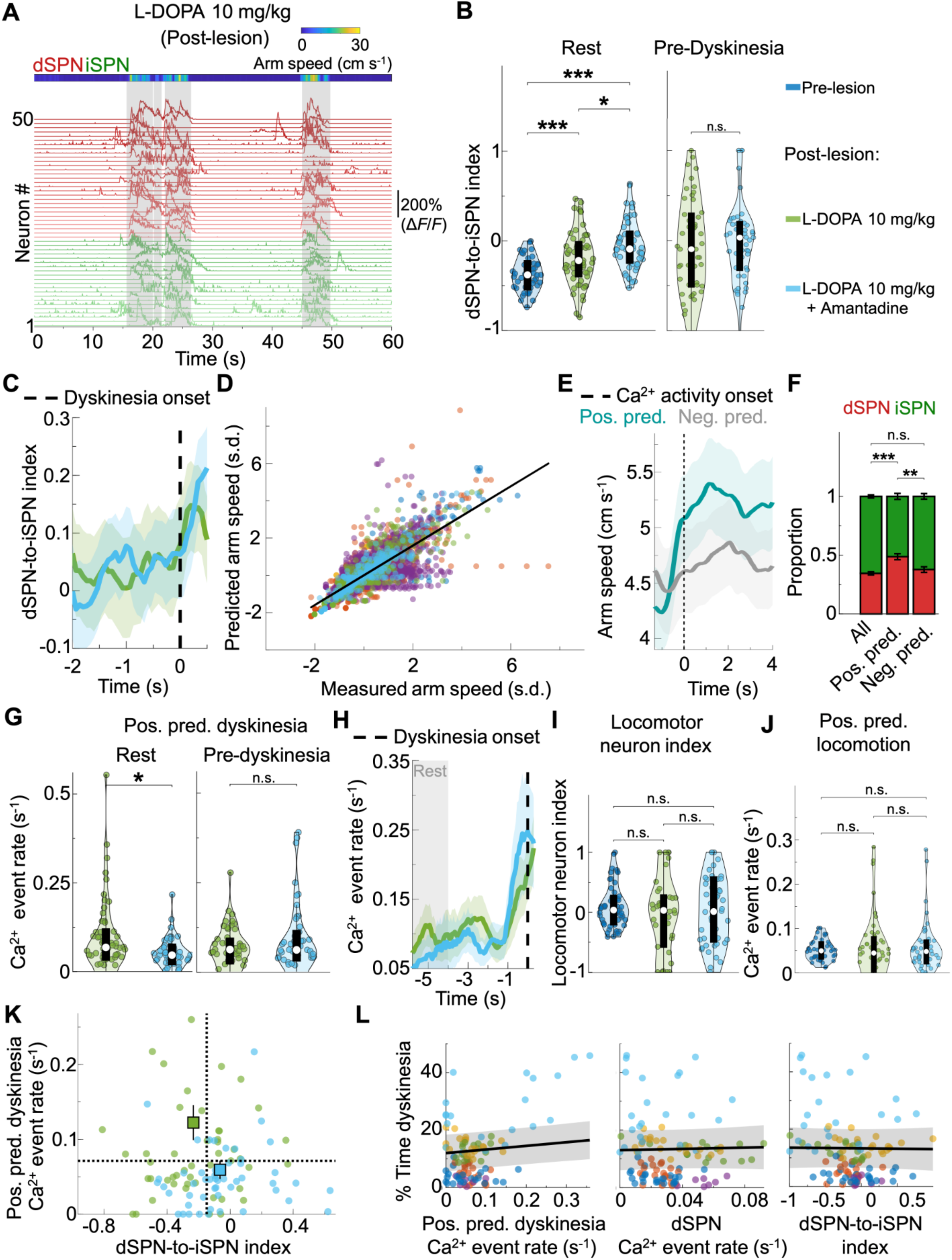
In the hyperkinetic condition, amantadine reduces the activity of dyskinesia-specific ensembles without improving dSPN to iSPN balance. (**A**) *Bottom*, Example Ca²⁺ activity traces (Δ*F*/*F*) of individual dSPNs (red) and iSPNs (green) recorded simultaneously during a L-DOPA (10 mg/kg) block. *Top*, Color plot of left forelimb speed during dyskinetic movements (marked with gray shading). **(B)** Violin plots of resting and pre-dyskinesia period dSPN-to-iSPN index values in individual experimental blocks across conditions. (***p < 0.001; n.s. = not significant; two-sided rank sum tests corrected for multiple comparisons using a Benjamini–Hochberg procedure with a false-discovery rate of 0.05; n = 56 pre-lesion blocks, 57 L-DOPA (10 mg/kg) blocks, and 55 L-DOPA (10 mg/kg) + amantadine blocks from 6 mice). **(C)** Mean dSPN-to-iSPN index values plotted relative to dyskinesia onset time (vertical dashed line) for each condition. Colored shading: s.e.m. for n = 57 L-DOPA (10 mg/kg) blocks, and 55 L-DOPA (10 mg/kg) + amantadine blocks from 6 mice. **(D)** Results of multivariate ridge regression analyses (for each mouse, imaging plane, and condition combination) predicting the speed of upcoming left forelimb movement, based on neural activity patterns just prior to dyskinetic movement. The scatter plot shows predicted *vs*. measured left forelimb speeds across 6 mice, color-coded by mouse. Each data point shows results for an individual dyskinetic movement. Speeds are z-scored for each mouse. Data is from L-DOPA (10 mg/kg) and L-DOPA (10 mg/kg) + amantadine conditions and pooled here for visualization purposes. The overall regression line (black) has a training-set R² of 0.59 and slope of 0.8. One outlier point (x = 1.65, y = 16.6) is not plotted to avoid visual distortion of the plot. **(E)** Mean speed of left forelimb movement relative to the time of Ca²⁺ events for positive-predictive and putative negative-predictive dyskinesia from the ridge regression model in (**D**), aggregated across all Ca²⁺ events. Although the activity of positive-predictive neurons (teal) was associated with an increase in dyskinesia forelimb speed, the activity of the putative negative-predictive neurons (gray) was not associated with a decrease in dyskinesia forelimb speed. Hence, subsequent analyses focused on positive-predictive dyskinesia neurons. Shading: s.e.m. for n = 111 experimental blocks, comprising 57 L-DOPA (10 mg/kg) blocks and 54 L-DOPA (10 mg/kg) + amantadine blocks from 6 mice. (One L-DOPA (10 mg/kg) + amantadine block excluded from here and in any subsequent analyses requiring identified dyskinesia-predictive neurons because none were identified in that block). Dashed vertical line: Ca^2+^ event onset (*t* = 0). **(F)** Proportions of dSPNs and iSPNs across the sets of all neurons, positive-predictive dyskinesia neurons, and negative-predictive dyskinesia neurons. (**p < 0.01, ***p < 0.001; n.s. = not significant; two-sided rank sum tests corrected for multiple comparisons using a Benjamini–Hochberg procedure with a false-discovery rate of 0.05; n = 111 experimental blocks, comprising 57 L-DOPA (10 mg/kg) blocks and 54 L-DOPA (10 mg/kg) + amantadine blocks from 6 mice). Error bars: s.e.m. **(G)** Violin plots of mean Ca^2+^ event rate during resting (*left*) and pre-dyskinesia (*right*) periods for positive-predictive dyskinesia neurons in individual experimental blocks. Outlier points > 3 s.d. from the mean not shown to avoid visual distortion of the plot. (*p < 0.05; n.s. = not significant; two-sided rank sum tests; n = 57 L-DOPA (10 mg/kg) blocks, 54 L-DOPA (10 mg/kg) + amantadine blocks from 6 mice). **(H)** Mean Ca^2+^ event rate of positive-predictive dyskinesia neurons plotted relative to dyskinesia onset time (vertical dashed line) for each condition. Gray shading indicates a time period classified as rest. Colored shading: s.e.m. for n = 57 L-DOPA (10 mg/kg) blocks, 54 L-DOPA (10 mg/kg) + amantadine blocks from 6 mice. **(I)** Violin plots of the mean resting period Ca^2+^ event rates for positive-predictive locomotion neurons per block across conditions. Outlier points > 3 s.d. from the mean not shown to avoid visual distortion of the plot. (n.s. = not significant; two-sided rank sum tests; n = 56 pre-lesion blocks, 57 L-DOPA (10 mg/kg) blocks, 54 L-DOPA (10 mg/kg) + amantadine blocks from 6 mice). **(J)** Violin plots of the mean locomotor neuron index (**Supp. Methods**) per block across conditions. (n.s. = not significant; two-sided rank sum tests; n = 56 pre-lesion blocks, 57 L-DOPA (10 mg/kg) blocks, 54 L-DOPA (10 mg/kg) + amantadine blocks from 6 mice) **(K)** Scatter plot of the mean resting period dSPN-to-iSPN index value (X-axis) versus positive-predictive dyskinesia neuron Ca^2+^ event rate (Y-axis). Each point represents a single experimental block, colored by condition (see legend in **B**). Large squares with error bars denote the mean ± s.e.m. for each condition. **(L)** Linear mixed-effects models showed that the resting activity of positive-predictive dyskinesia neurons significantly predicted the percentage of time spent in dyskinesia per block (*left*, slope = 12.6, p = 0.002). By contrast, resting dSPN activity (*middle*) and the resting dSPN-to-iSPN index (*right*) were not significant predictors in the same analysis. Scatter plots show individual blocks, color-coded by individual mice, with regression lines fit for the L-DOPA (10 mg/kg) condition. The x-axis is clipped at 3 s.d. from the mean to avoid visual distortion of the plot owing to a small number (1 to 3 per plot) of outlier data points (n = 111 blocks from 6 mice).

We therefore hypothesized that amantadine, as in the hypokinetic state, may instead act on action-specific neural ensembles.^9,34,41^ We first identified a forelimb dyskinesia-predictive ensemble using a multivariate ridge regression model. In the training data, neural activity during the 4-s window preceding dyskinesia onset predicted the speed of the upcoming dyskinetic movement with a training-set R² of 0.59 (**Figures 3D, S9B**,**C**). This model identified a population of positive-predictive dyskinesia neurons whose Ca^2+^ events were associated with a significant increase in forelimb speed (**Supp. Methods**) (**Figure 3E**). We did not find a functionally validated negative-predictive population; therefore, subsequent analyses focused on the positive-predictive ensemble. This dyskinesia-predictive ensemble was composed of both dSPNs and iSPNs, but was relatively enriched for dSPNs compared to the overall detected-neuron population (∼49% dSPN vs. ∼35% overall; **Figure 3F**).^41^ With high-dose L-DOPA treatment, both iSPNs and dSPNs showed heterogeneity in their responses across ensemble classes. In both pathways, neurons outside the locomotion and dyskinesia ensembles were suppressed relative to the pre-lesion baseline population, whereas neurons within the forelimb dyskinesia ensemble showed significantly elevated activity relative to both baseline and the non-ensemble population (**Figure S10A**).

Amantadine treatment selectively reduced the resting Ca^2+^ event rate of the positive-predictive dyskinesia ensemble, without a detectable effect during the immediate pre-dyskinesia period or the first second of dyskinetic behavior (**Figures 3G,H, Figure S6B**). This action appeared selective as amantadine had no effect on the resting activity of the positive-predictive locomotion neurons or the overall locomotor neuron index (**Figures 3I,J**). This corresponds with our behavioral data showing that amantadine reduced dyskinesia without impacting locomotion (**Figure S11A**).

We next plotted amantadine’s effects on pathway balance and ensemble activity in the dyskinetic condition: the treatment reduced the activity of the dyskinesia ensemble without improving pathway balance (**Figure 3K**).

Finally, linear mixed-effects models supported the functional relevance of the positive-predictive dyskinesia ensemble, as its resting activity significantly predicted the total time spent exhibiting dyskinetic movements per block. By contrast, the resting dSPN-to-iSPN index, dSPN activity, and iSPN activity did not (**Figure 3L**).

### The Ensembles Predicting Locomotion and Dyskinesia Are Distinct Neural Populations

To compare the neural representations of locomotion and forelimb dyskinetic movements, we focused on experimental time blocks that had separate bouts of both locomotion and dyskinesia. Like prior sections, we used a multivariate ridge regression model to identify positive-predictive locomotion neurons and positive-predictive dyskinesia neurons. Visually, these two ensembles appeared to be spatially distinct (**Figure 4A**). We then confirmed this functional separation by examining their activity aligned to movement onset. As expected, the locomotion predictive ensemble was more active preceding locomotion, while the dyskinesia predictive ensemble was more active preceding dyskinesia (**Figure 4B**). Additionally, across the population of SPNs, regression coefficients for dyskinesia and locomotion predictive cells were largely uncorrelated (R^2^ ranged from 0.01 to 0.07 across mice; **Figure 4C**). Although we independently classified neurons as positive locomotion and positive dyskinesia predictive, only a limited subset of cells had both classifications (9.3 ± 10.5%; mean ± s.d.; n = 41 blocks with L-DOPA (10 mg/kg) treatment with both locomotion and dyskinesia bouts; 519 positive-predictive locomotion and/or dyskinesia neurons across 6 mice).

**Figure 4.**
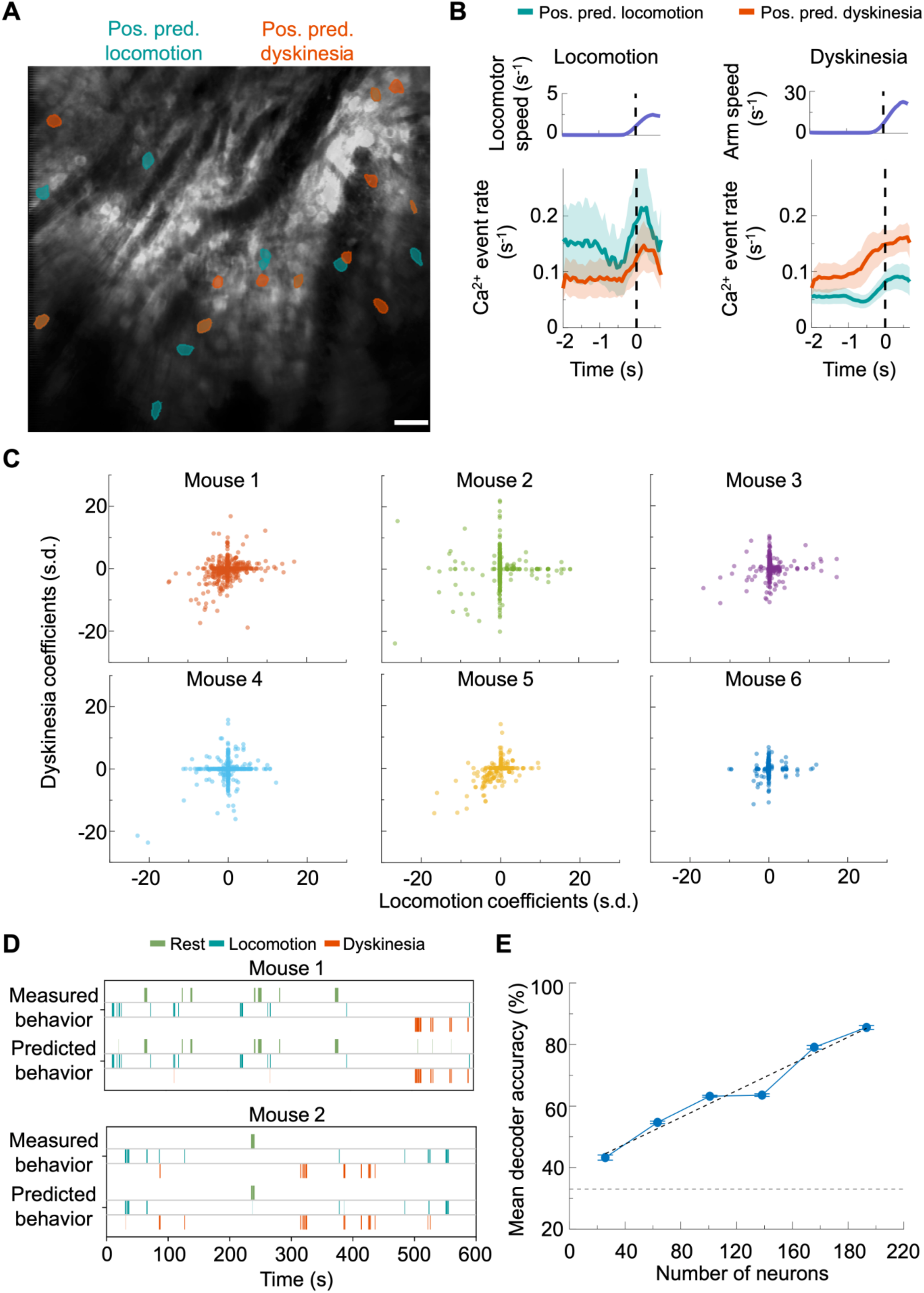
Distinct striatal neural ensembles encode locomotion and dyskinesia. (**A**) Example fluorescence image of striatal neurons visualized by two-photon microscopy via a GRIN lens. Masks for positive-predictive locomotion and dyskinesia neurons are overlaid on the image to illustrate their spatial distribution. The image is a time-averaged projection of a 10 min Ca^2+^ video taken in the L-DOPA (10 mg/kg) condition. Scale bar: 40 µm. **(B)** *Lower,* Mean Ca^2+^ event rates of positive-predictive locomotion and dyskinesia neurons, aligned to locomotion (*left*) or dyskinesia (*right*) onset during L-DOPA (10 mg/kg) blocks. *Upper*, plots of mean locomotion speed or dyskinetic arm speed. Vertical dashed lines mark the times of movement onset and offset. Shading: s.e.m. for n = 41 blocks from 6 mice. **(C)** Scatter plot showing normalized locomotion and normalized dyskinesia coefficients across 6 individual mice. Each point denotes data from a single neuron. The low R² values across mice (Mouse 1: 0.06, Mouse 2: 0.01, Mouse 3: 0.02, Mouse 4: 0.07, Mouse 5: 0.33, Mouse 6: 0.01) indicate that locomotion– and dyskinesia-predictive neurons are largely distinct populations. **(D)** Example time series from two experimental blocks, showing empirically measured behavioral states (top rows) and predictions from an SVM decoder trained on striatal neural activity (bottom rows) (**Supp. Methods**). Each colored segment represents a period classified as rest (green), locomotion (cyan), or dyskinesia (orange). For these blocks, the decoder successfully distinguished rest, locomotion, and dyskinesia states (95% and 90% accuracy on a held-out test set for the blocks shown for mouse 1 and mouse 2, respectively). **(E)** Mean classification accuracy of the SVM decoders as a function of the number of cells recorded per block, grouped into six bins. The positive trend (*R*² = 0.96, dashed black line) shows that with enough neurons recorded, behavioral states are distinguishable based on their accompanying neural activity patterns (dashed gray horizontal line; chance level of 33.2% ± 5.0%, as determined by training on shuffled labels; n = 112 L-DOPA (10 mg/kg) and L-DOPA (10 mg/kg) + amantadine blocks from 6 mice). Error bars: s.e.m.

To determine whether locomotion and dyskinesia are represented as distinct neural states, we created support vector machine (SVM) decoders to predict the animals’ motor behavioral states based on their SPN activity patterns. Time series visualizations of decoder outputs demonstrated their capabilities to predict periods of locomotion, dyskinesia, and rest (**Figures 4D**, **S12A**), especially when the datasets included large numbers of neurons (**Figure 4E**). Altogether, these findings argue that the dorsolateral striatum has distinct SPN ensemble representations of locomotion and forelimb dyskinesia.

## DISCUSSION

The striatum plays a central role in motor control, integrating inputs from cortical, thalamic, and dopaminergic sources to guide actions.^42^ While the classical view of the striatum emphasizes opposing dSPN/iSPN roles^1,11^, normal motor control often requires coordinated, action-specific activity in both pathways.^16–19^ Our study supports a model in which striatal pathophysiology and its treatment can be understood along at least two distinct mechanisms: the regulation of pathway balance and the modulation of action-specific ensembles (**Figure 5**).

**Figure 5.**
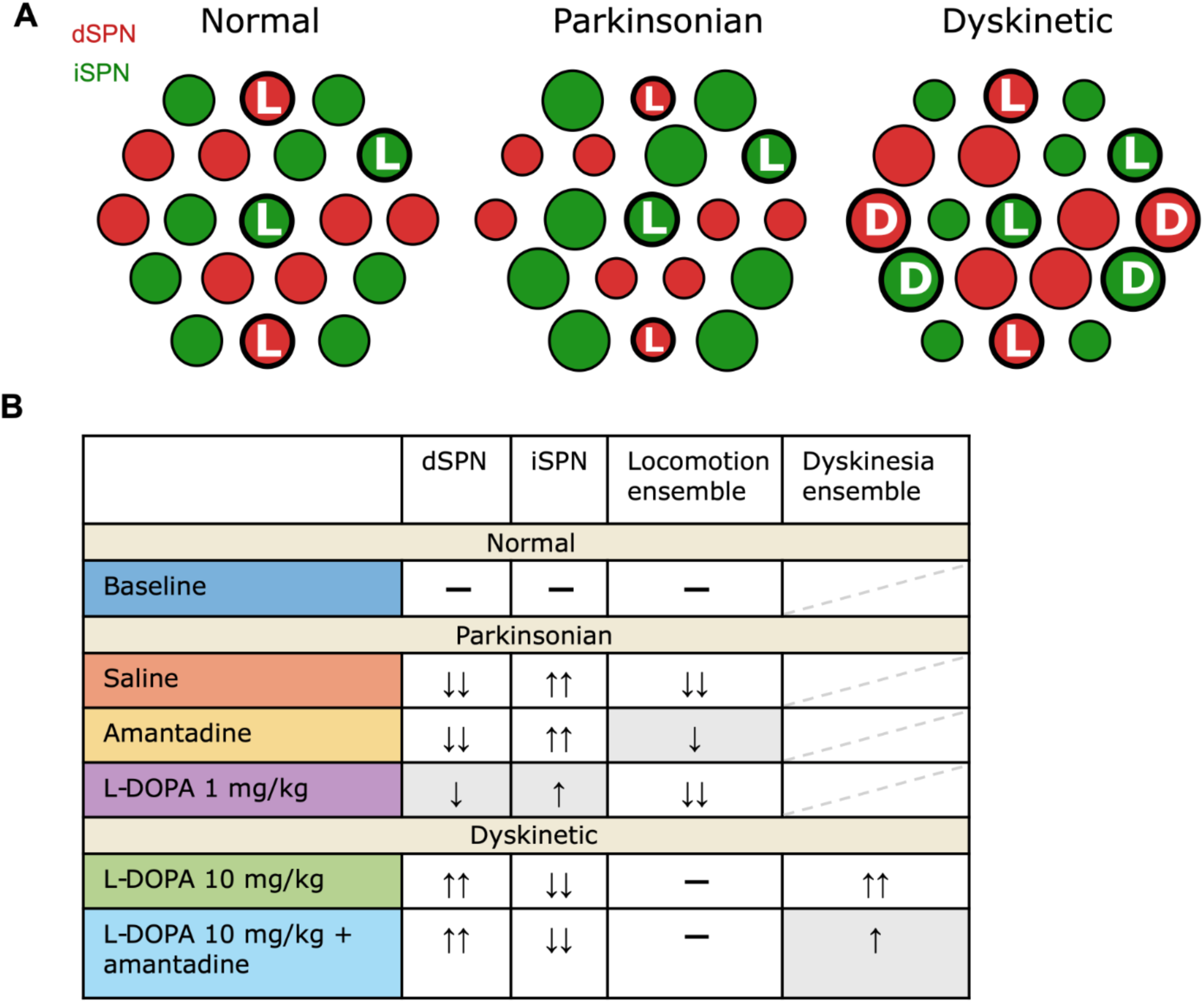
Dissociable therapeutic mechanisms of L-DOPA and amantadine. (**A**) Schematic illustrating two functional properties of SPNs: pathway identity and action-specific ensemble membership. Red circles represent dSPNs, and green circles represent iSPNs. Circle size repreesents relative activity level, with smaller circles indicating reduced activity and larger circles indicating increased activity. Thick black borders denote neurons belonging to action-specific ensembles. **L** marks locomotion-predictive ensemble neurons, and **D** marks forelimb dyskinesia-predictive ensemble neurons. In the normal state, dSPN and iSPN activity are relatively balanced, with locomotion-predictive neurons distributed across both pathways. In the parkinsonian state, pathway balance shifts toward iSPNs and locomotion-ensemble activity is reduced. In the dyskinetic state, pathway balance shifts toward dSPNs and a distinct forelimb dyskinesia-predictive ensemble is present. **(B)** Summary table of pathway– and ensemble-level activity changes across conditions. Arrows indicate activity changes relative to the pre-lesion baseline condition; double arrows indicate larger changes than single arrows, and dashes indicate no major detected change. Gray shading highlights the primary therapeutic effect emphasized for each treatment. In saline-treated parkinsonian mice, dSPN/iSPN balance and locomotion-ensemble activity were reduced (Figures 2A**–B****, 2F–H**). Amantadine partially restored locomotion-ensemble activity without restoring pathway balance (Figures 2A**–B****, 2F–H**). Low-dose L-DOPA partially restored pathway balance but not locomotion-ensemble activity (Figures 2A**–B****, 2F–H**). In the dyskinetic condition, high-dose L-DOPA shifted pathway balance toward dSPNs and increased activity in the forelimb dyskinesia-predictive ensemble (Figures 3B**,G****,H,K**). Adding amantadine suppressed the resting activity of the forelimb dyskinesia ensemble without restoring pathway balance or altering locomotion-coding ensembles (Figures 3B**,G****–K**). Together, these findings support a model in which L-DOPA primarily acts through pathway rebalancing, whereas amantadine preferentially modulates action-specific ensemble activity.

### Action-Specific Ensembles Provide a Complementary Framework to Pathway Balance

In the hypokinetic state we observed the classical shift in pathway balance toward iSPNs and a disruption in locomotion-predictive ensemble activity. Similarly, in the hyperkinetic state, in addition to a classical shift in pathway balance towards dSPNs, we identified a distinct forelimb dyskinesia-predictive ensemble. The ensemble-based view of dyskinetic movement is consistent with work from Girasole et al., who established that a specific and stable subpopulation of striatal neurons is required for levodopa-induced dyskinesias.^9^ This result was recently extended by Alcacer et al., who found that different types of dyskinetic movement are encoded by distinct ensembles of dSPNs and iSPNs.^41^

The resting activity of these ensembles predicted behavioral measures across blocks, whereas the overall dSPN-to-iSPN imbalance did not. This suggests that while pathway balance may help establish a permissive state for movement, the specific instruction to move may depend on ensemble coding activity.

Our findings also suggest therapeutic relevance for these action-coding ensembles. Amantadine improved behavior in both states without restoring pathway balance, and its clearest neural effect was selective modulation of action-specific ensembles during resting periods, with little detectable effect during the immediate pre-movement period or early movement execution. These findings suggest that amantadine may exert its behavioral effects in part by modulating pre-action state-setting mechanisms. This is consistent with our finding that action-specific predictive signals emerged during resting periods preceding movement onset and recent work identifying the striatum as an important contributor to the timing of upcoming actions.^43^ However, our data are correlational and do not establish that the ensemble changes are sufficient or necessary for amantadine’s behavioral effects.

### A Novel Approach for Therapeutic Development Revealed by Amantadine

Modulating action-specific neural activity may offer a therapeutic alternative to pathway rebalancing. Our combined imaging and behavioral assay could screen for interventions that normalize pathological ensemble activity.

Amantadine’s pharmacology could provide a starting point for this approach.^21,22^ Although our study did not determine which of amantadine’s targets mediate its ensemble-specific effects, our results allow us to generate hypotheses. Two targets of interest are NMDA receptors and Kir2 potassium channels.^21,26,27^ NMDA receptors have an important role in corticostriatal plasticity, which seems to be impaired in parkinsonian and dyskinetic brain states.^44–47^ Amantadine’s NMDA receptor antagonism might limit long-term potentiation (LTP) at corticostriatal synapses onto action-specific SPN populations, potentially reducing maladaptive plasticity and pathological activity in these neurons.^21,27^ Amantadine’s antagonism of Kir2 receptors may also alter SPN excitability and plasticity in a pathway-dependent manner.^26^ Future studies could test whether modifying amantadine or related compounds to bias effects toward particular receptor classes improves behavioral outcomes while normalizing striatal ensemble dynamics.^48^

### Limitations of the Study

Our study has several limitations. First, it only examined one hypokinetic condition (severe acute unilateral dopamine depletion via 6-OHDA) and one hyperkinetic condition with one dyskinetic movement (L-DOPA-induced forelimb dyskinesias in unilateral dopamine-depleted mice). Our results may not generalize to other hypokinetic or hyperkinetic conditions or additional types of L-DOPA-induced dyskinesias (e.g. orolingual dyskinesias or axial dyskinesias). Second, although we identified neural populations with activity that correlated with specific behaviors, we did not directly manipulate the activity of these populations. Thus, we cannot establish definitive causal links between their activity patterns and observed behavioral changes. However, the fact that resting activity reliably predicted upcoming movement characteristics supports these ensembles’ role in action specification. Third, we did not track individual neurons longitudinally, instead identifying action-specific SPNs separately for each condition. This limited our ability to determine whether locomotion– or dyskinesia-predictive ensembles identified in the absence of amantadine were the same ensembles amantadine subsequently modulated. Exploratory analyses of temporally proximal sessions suggested some degree of overlap in positive-predictive ensembles across nearby conditions, but this does not establish a stable longitudinal identity across the broader experiment. Fourth, because fewer dSPNs than iSPNs had detectable calcium activity in this preparation, the pathway-balance analyses should be interpreted as applying to the detected dSPN population available here rather than the full direct-pathway population. Although we used bootstrap subsampling to equalize detected cell counts, this procedure could not recover activity from undetected dSPNs. Fifth, we utilized a head-fixed behavioral preparation. While this constrained the behavioral repertoire and limited assessment of axial and some other dyskinetic subtypes, the setup still allowed us to examine core relevant pathophysiology; we observed robust hypokinetic and forelimb dyskinetic phenotypes and replicated established dSPN/iSPN activity imbalances. Accordingly, this paradigm is best viewed as a controlled, imaging-compatible assay of specific locomotor and dyskinetic components, rather than as a full behavioral recapitulation of freely moving parkinsonian and L-DOPA-induced dyskinetic phenotypes. Sixth, the mixed-effects analyses were based on only 5–6 animals. Although the number of imaging blocks per condition was substantial, the mouse random effect was therefore estimated from a modest number of levels, and these results should be interpreted cautiously.

Future studies should examine additional movement disorder models, extend ensemble analyses to freely moving behavior, manipulate specific neural ensembles, and track individual SPNs across conditions.

## Author Roles

G.C. and M.J.S. conceived the study and designed data analyses. G.C. performed behavioral and imaging experiments, surgeries, and data analyses. R.C. built the microscope and created the software to control the microscopy and behavioral apparatus. J.L. bred transgenic mice and performed immunohistochemistry. Y.Z. designed and performed viral vector subcloning and virus validation. F. D. developed key software used in the data analyses for this study. G.C. and M.J.S. wrote the manuscript. All authors edited the paper. M.J.S. supervised the study.

## Acknowledgements

G.C. thanks B. Ahanonu, T. Rogerson, and O. Hazon for technical advice regarding rodent surgeries and data analyses.

## Financial Disclosures of All Authors (for the Past 12 Months)

The authors report no disclosures.

## RESOURCE AVAILABILITY

### Lead contact

Further information and requests for resources should be directed to lead contact Mark J. Schnitzer.

### Materials availability

This study did not generate new unique reagents.

### Data and code availability

All data and code used in this paper are available from the lead contact upon request.

**Supplemental Figure 1.**
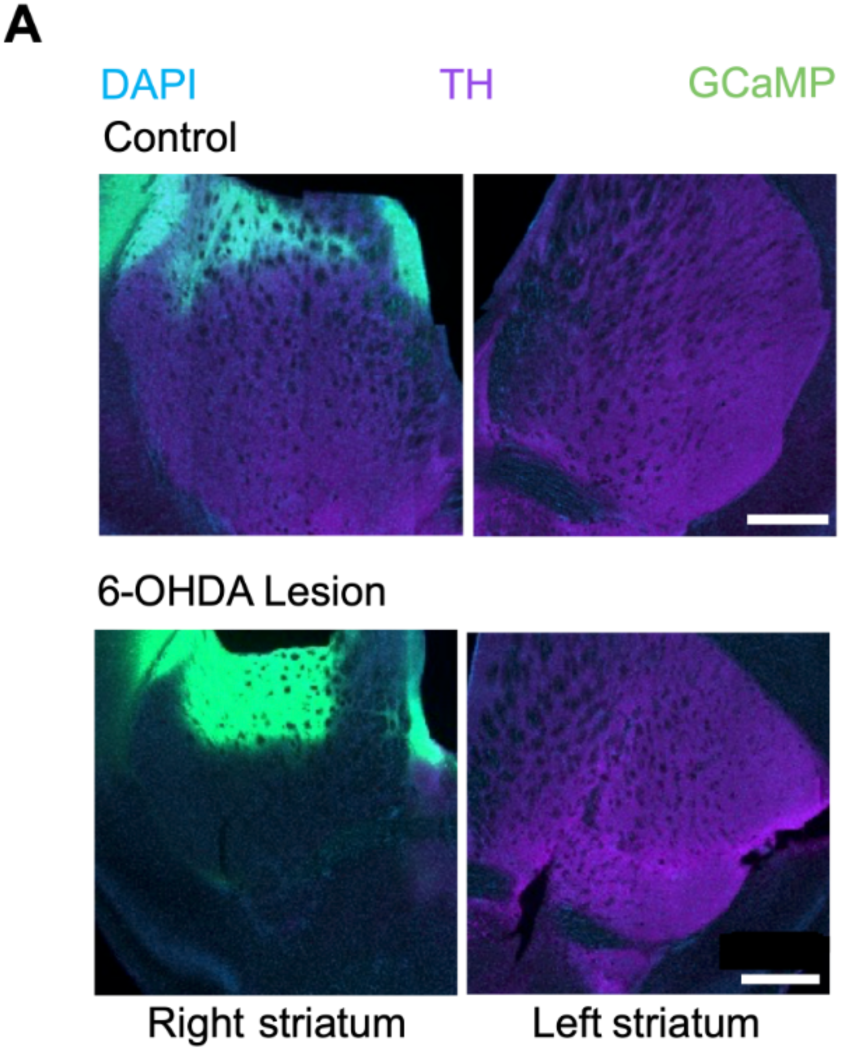
Validation of dopamine depletion via fluorescence immunostaining for tyrosine hydroxylase. (**A**) We immunostained brain slices for tyrosine hydroxylase to evaluate the loss of dopaminergic axons in the striatum (Purple = Tyrosine Hydroxylase (TH) immunostaining; green = jGCaMP7f; blue = DAPI). *Top*, Images of tissue slices from an example control mouse, showing relatively symmetric tyrosine hydroxylase staining bilaterally. *Bottom*, Images of tissue slices from a mouse with a unilateral 6-OHDA lesion of the SNc, showing loss of dopaminergic axons in the right striatum, ipsilateral to the lesioned SNc. Scale bars: 500 µm.

**Supplemental Figure 2.**
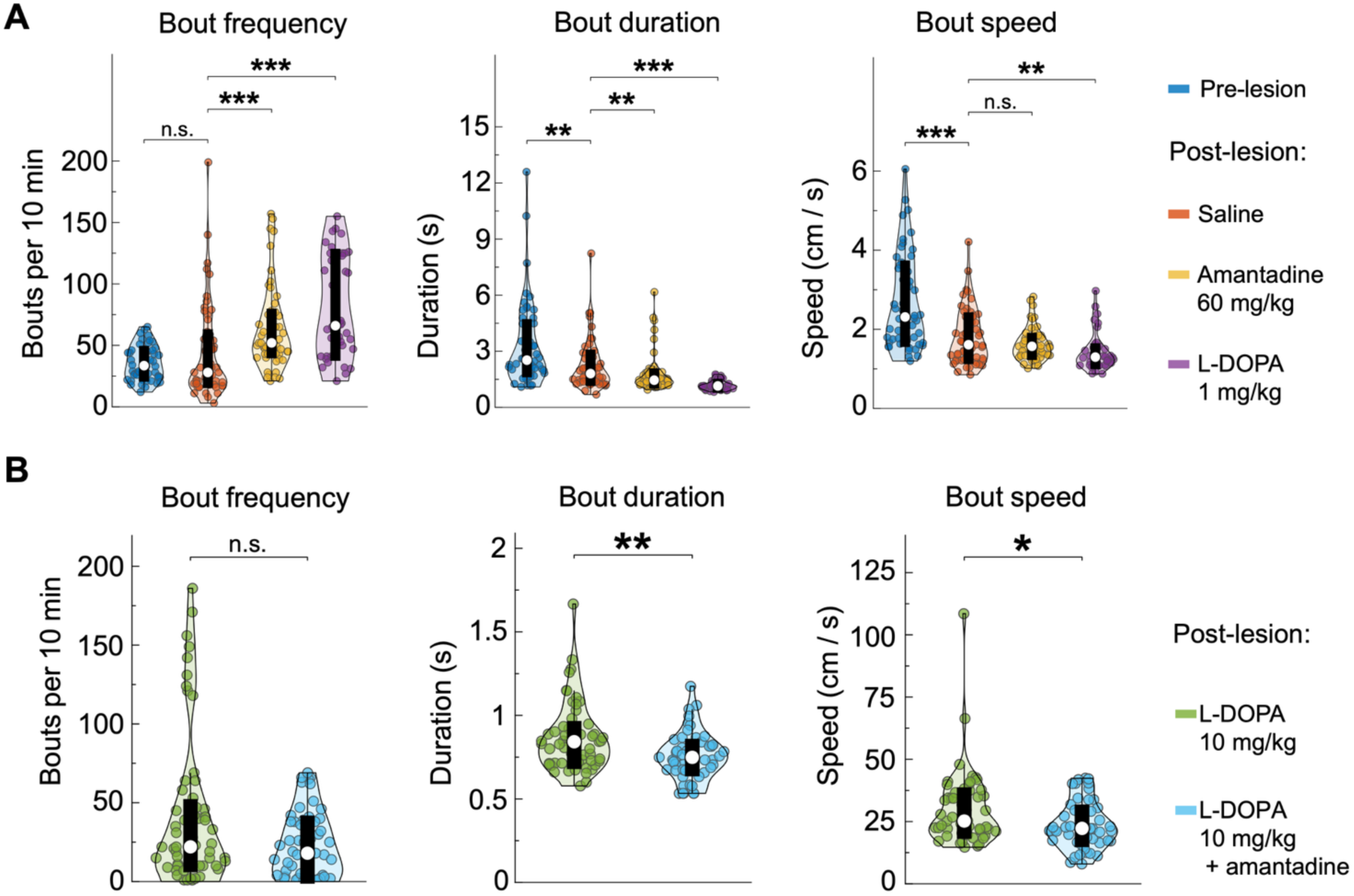
L-DOPA and amantadine increase locomotion bout frequency, whereas amantadine reduces dyskinetic bout duration and speed. (**A**) Violin plots showing per-block locomotion bout frequency (bouts per 10-min block), mean bout duration, and mean bout speed across hypokinetic conditions. (**p < 0.01, ***p < 0.001; n.s. = not significant; two-sided rank sum tests, corrected for multiple comparisons using a Benjamini–Hochberg procedure with a false-discovery rate of 0.05; n = 50 pre-lesion blocks, 53 saline blocks, 53 amantadine blocks, and 38 L-DOPA (1 mg/kg) blocks from 5 mice). **(B)** Violin plots showing per-block dyskinetic bout frequency (bouts per 10-min block), mean forelimb dyskinesia bout duration, and mean forelimb dyskinesia bout speed across hyperkinetic conditions. (*p < 0.05; two-sided rank sum tests, corrected for multiple comparisons using a Benjamini–Hochberg procedure with a false-discovery rate of 0.05; n = 56 pre-lesion blocks, 57 L-DOPA (10 mg/kg) blocks, and 56 L-DOPA (10 mg/kg) + amantadine blocks from 6 mice).

**Supplemental Figure 3.**
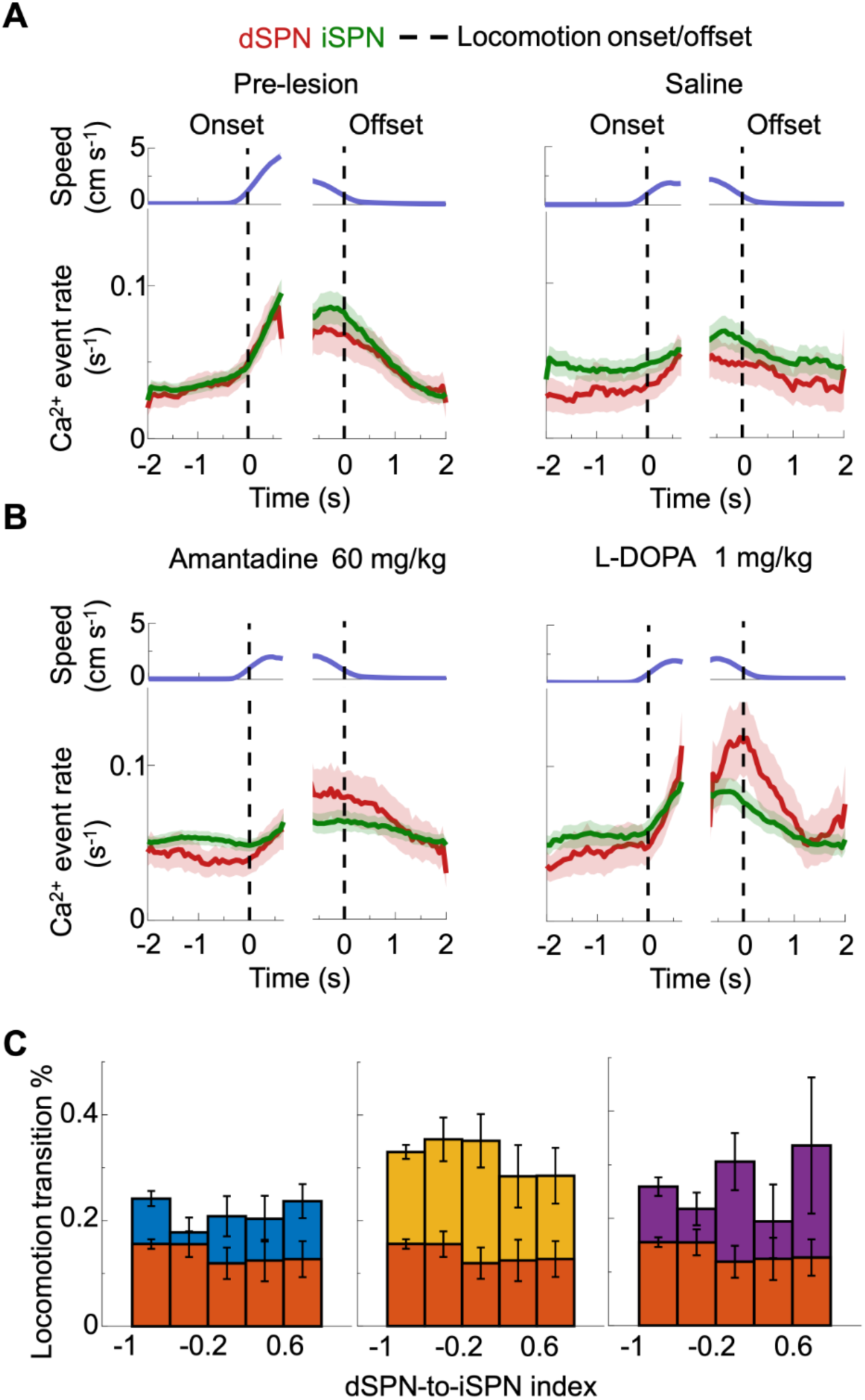
Locomotion-related SPN dynamics and locomotion initiation probabilities across conditions. (**A**) *Top,* Mean locomotor speed, plotted relative to times of locomotor onset and offset (vertical dashed lines), before (*left*) or after (*right*) unilateral dopamine lesion of the SNc. *Bottom*, Ca²⁺ event rates of dSPNs and iSPNs plotted relative to locomotion onset and offset, shown for each condition. Colored shading: s.e.m. for n = 50 pre-lesion and 53 saline blocks from 5 mice. **(B)** Analogous plots to those of A, for mice administered either amantadine or L-DOPA (1 mg/kg) after the dopamine lesion. Colored shading: s.e.m. for n = 53 amantadine blocks and 38 L-DOPA blocks, from 5 mice. **(C)** Bar plots showing the probabilities of transitioning from rest to locomotion as a function of the dSPN-to-iSPN index in mice at rest. Pre-lesion, amantadine, and L-DOPA (1 mg/kg) conditions displayed significantly higher transition probabilities across all index bins as compared to the saline condition. Error bars: s.e.m. estimated based on counting statistics for a binomial distribution (**Supp. Methods**).

**Supplemental Figure 4.**
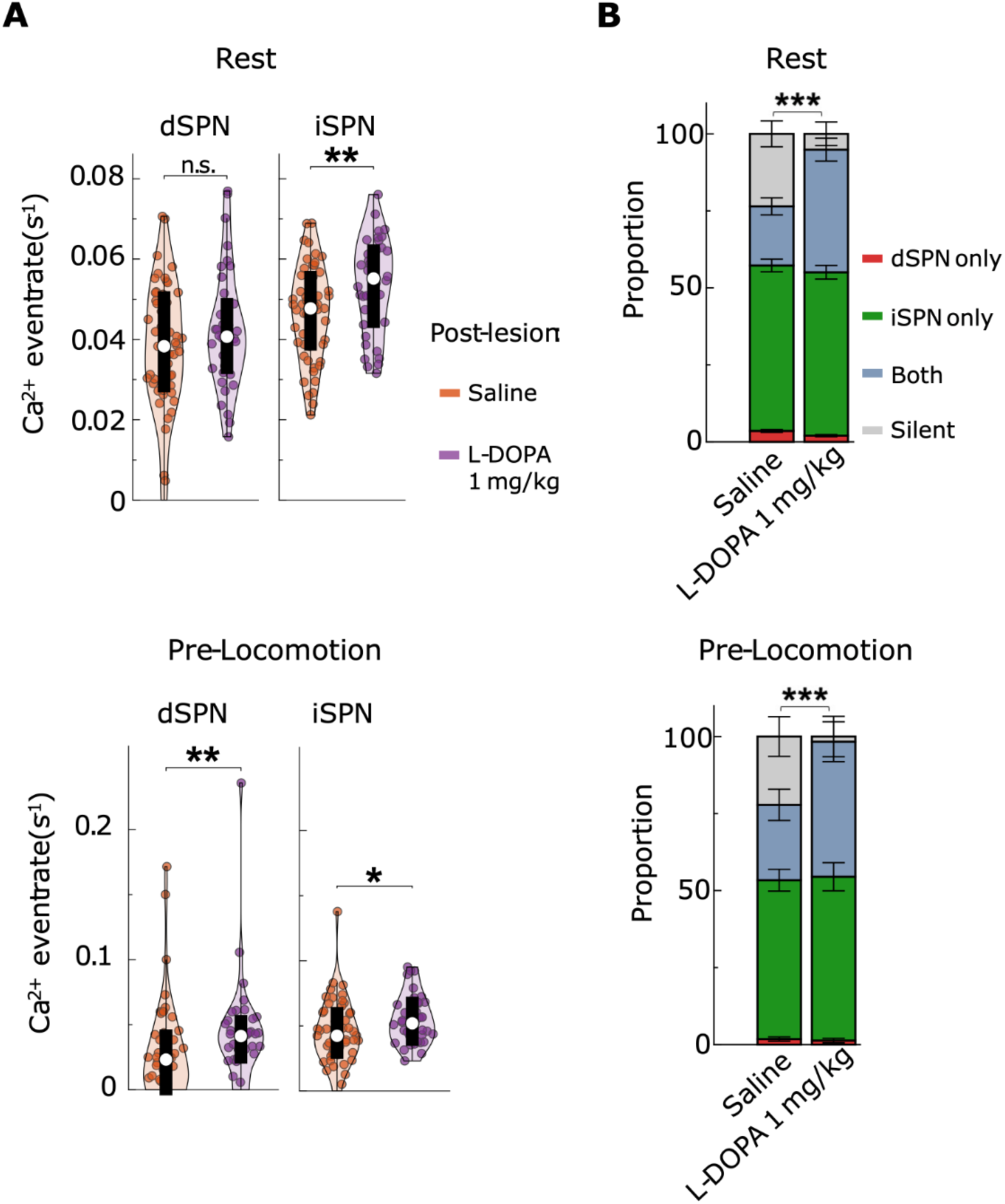
Low-dose L-DOPA (1 mg/kg) increases dSPN participation in the parkinsonian condition. (**A**) Violin plots of resting period (*top*) and pre-locomotion period (*bottom*) mean Ca^2+^ event rates of dSPN and iSPN neurons neurons in the parkinsonian condition with saline control injection vs. low-dose L-DOPA (1 mg/kg) injection. (*p < 0.05; **p < 0.01; n.s. = not significant; two-sided rank sum tests; n = 53 saline blocks and 38 L-DOPA (1 mg/kg) blocks from 5 mice). **(B)** Stacked bar plots of resting period (*top*) and pre-locomotion period (*bottom*) showing proportion of periods with dSPN activity only, iSPN activity only, activity from both dSPNs and iSPNs, and neither (silent) in the parkinsonian condition with saline control injection vs. low-dose L-DOPA (1 mg/kg) injection. dSPN participation (defined as proportion of periods per block with detected dSPN activity) was statistically significantly higher for the L-DOPA (1 mg/kg) injected condition vs. saline. (***p < 0.001; two-sided rank sum tests; n = 53 saline blocks and 38 L-DOPA (1 mg/kg) blocks from 5 mice). Error bars: s.e.m.

**Supplemental Figure 5.**
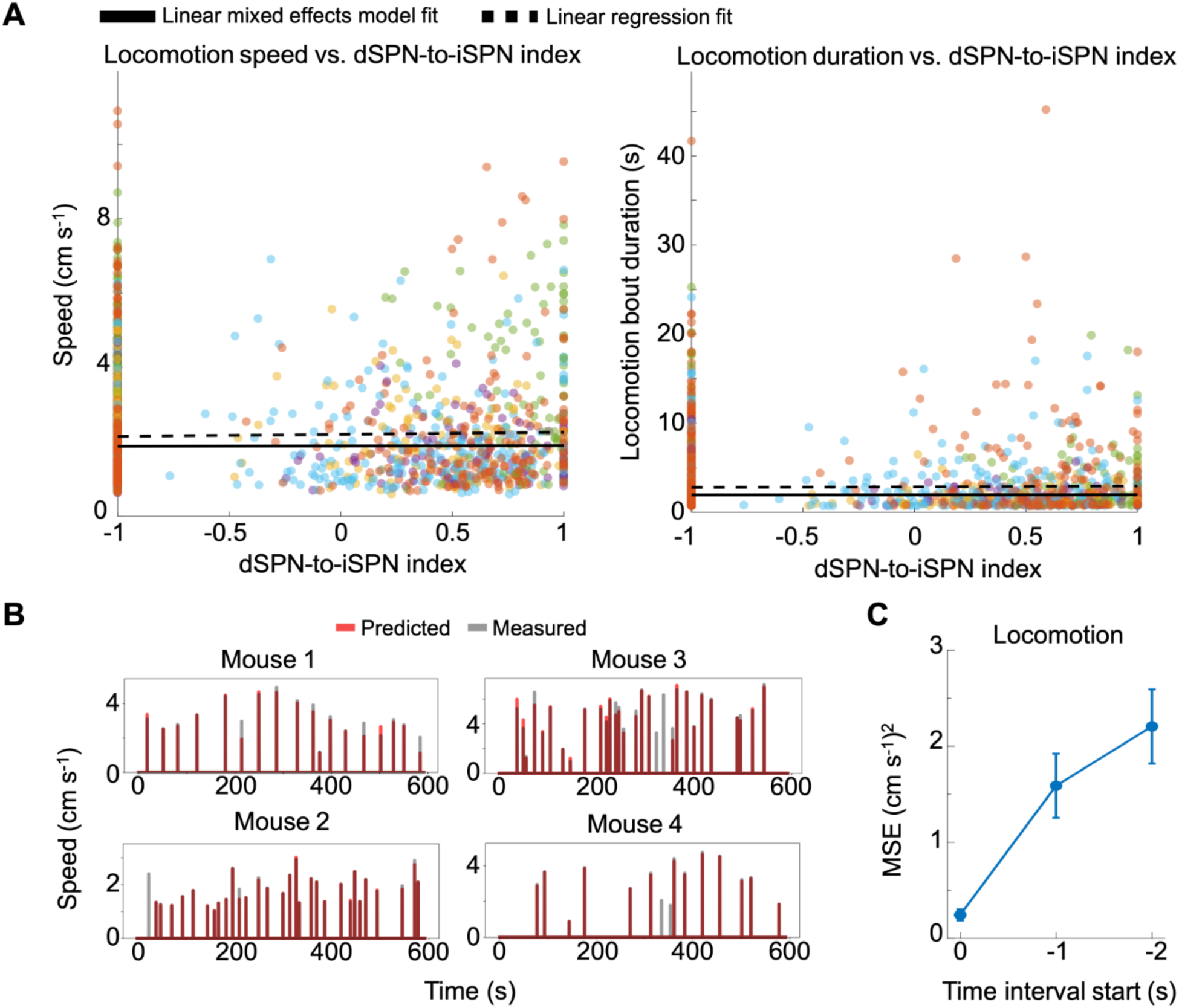
Neural ensemble representations, but not the dSPN-to-iSPN index, predict the speed of upcoming locomotion. (**A**) Scatter plots showing a lack of association between the dSPN-to-iSPN index during 2-s-long pre-locomotion periods and the speed (*left*) or duration (*right*) of the upcoming locomotion bout. Each data point denotes an individual locomotion bout, with colors denoting individual mice. Data is pooled across 50 pre-lesion, 53 saline, 53 amantadine, and 38 L-DOPA (1 mg/kg) blocks. Solid black lines: Fits from a linear mixed-effects model, in which the systematic variations across mice were treated as random effects and the variations across brain state and treatment conditions were treated as fixed effects. Dashed black lines: Fits from a least-squares regression applied to the pooled dataset without accounting for condition– or mouse-specific effects. For plots of locomotion speed *vs*. dSPN-to-iSPN index, the mixed-effects model yielded p = 0.757 and the linear regression yielded an R² < 0.001. For plots of locomotion bout duration *vs*. dSPN-to-iSPN index, the mixed-effects model yielded p = 0.937 and the linear regression yielded an R² < 0.001. **(B)** Example time series plots showing predicted (red) and empirically measured (gray) locomotor speeds right after locomotion onset, using the multivariate ridge regression model from **Figure 2C**. **(C)** Multivariate ridge regression performance results (**Figure 2C**, **Supp. Methods**) across different time intervals of 4-s-duration, shifted progressively further back in time from locomotion onset. The mean squared error (MSE) is plotted as a function of the start time of the 4-s-window, showing how the ability to predict locomotor speed based on neural activity patterns declines with increasing temporal separation from locomotion onset. Error bars: s.e.m. for n = 193 experimental blocks comprising 50 pre-lesion blocks, 53 saline blocks, 53 amantadine blocks, and 37 L-DOPA (1 mg/kg) blocks from 5 mice.

**Supplemental Figure 6.**
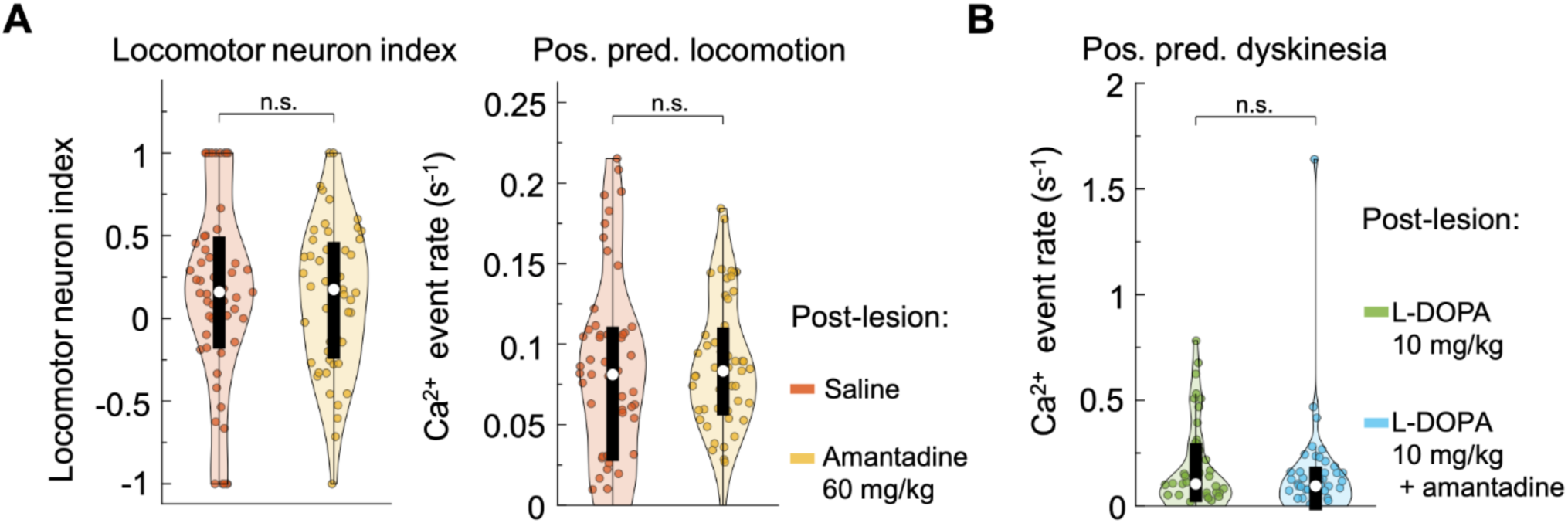
Amantadine does not significantly affect ensemble activity during behavior execution. (**A**) Violin plots showing the locomotor neuron index and mean Ca^2+^ event rate of positive-predictive locomotion neurons during the initial locomotion period (0 to 1 s) in the parkinsonian condition with amantadine injection vs. saline control injection. (n.s. = not significant; two-sided rank sum tests; n = 53 saline blocks and 53 amantadine blocks from 5 mice). **B)** Violin plots showing the mean Ca^2+^ event rate of positive-predictive dyskinesia neurons during the initial dyskinesia period (0 to 1 s) in the high-dose L-DOPA condition vs. high-dose L-DOPA + amantadine condition. (n.s. = not significant; two-sided rank sum tests; n = 57 L-DOPA (10 mg/kg) blocks, 54 L-DOPA (10 mg/kg) + amantadine blocks from 6 mice).

**Supplemental Figure 7.**
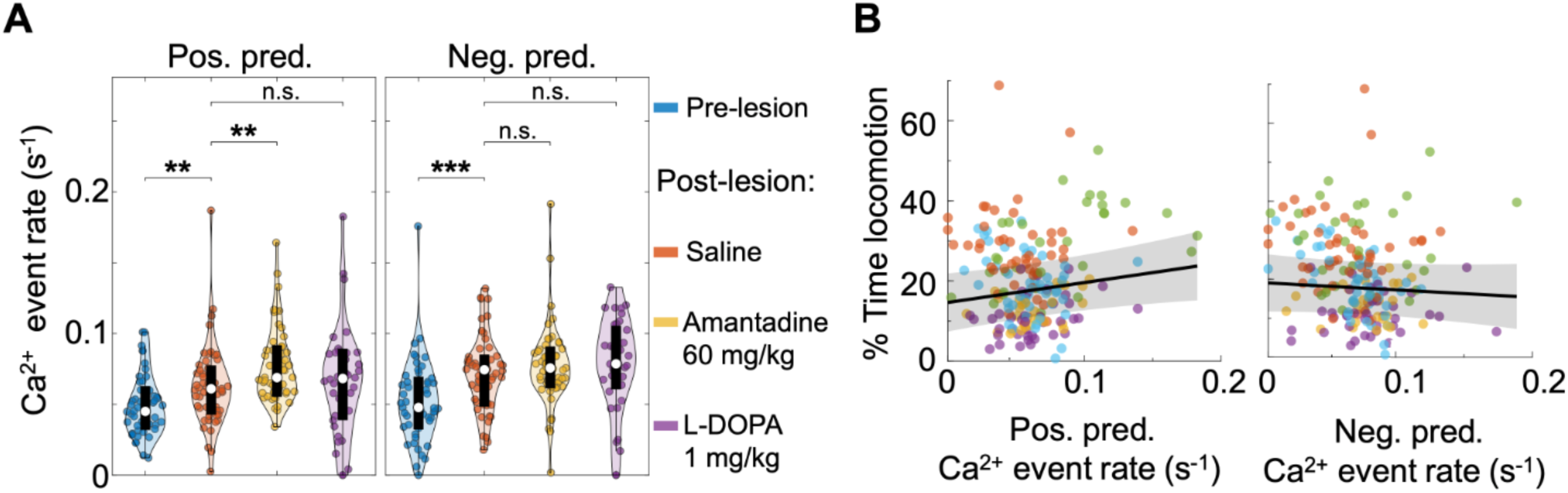
The resting activity of positive-predictive locomotion neurons increases with amantadine treatment and is predictive of the percentage of time spent in locomotion. (**A**) Violin plots of mean resting period Ca^2+^ event rates of positive– and negative-predictive locomotion neurons across conditions (**p < 0.01, ***p < 0.001; n.s. = not significant; two-sided rank sum tests corrected for multiple comparisons using a Benjamini–Hochberg procedure with a false-discovery rate of 0.05; n = 50 pre-lesion blocks, 53 saline blocks, 53 amantadine blocks, and 37 L-DOPA (1 mg/kg) blocks from 5 mice). **(B)** Linear mixed-effects models showed that the mean resting Ca^2+^ event rate of positive-predictive locomotion neurons significantly predicted the percentage of time spent locomoting per block (p = 0.03), whereas the mean resting Ca^2+^ event rate of negative-predictive locomotion neurons was not a significant predictor in the same analysis. Points denote data from individual experimental blocks, color-coded by mouse, with regression lines fit for the untreated hypokinetic (saline control) condition (n = 193 experimental blocks comprising 50 pre-lesion blocks, 53 saline blocks, 53 amantadine blocks, and 37 L-DOPA (1 mg/kg) blocks from 5 mice).

**Supplemental Figure 8.**
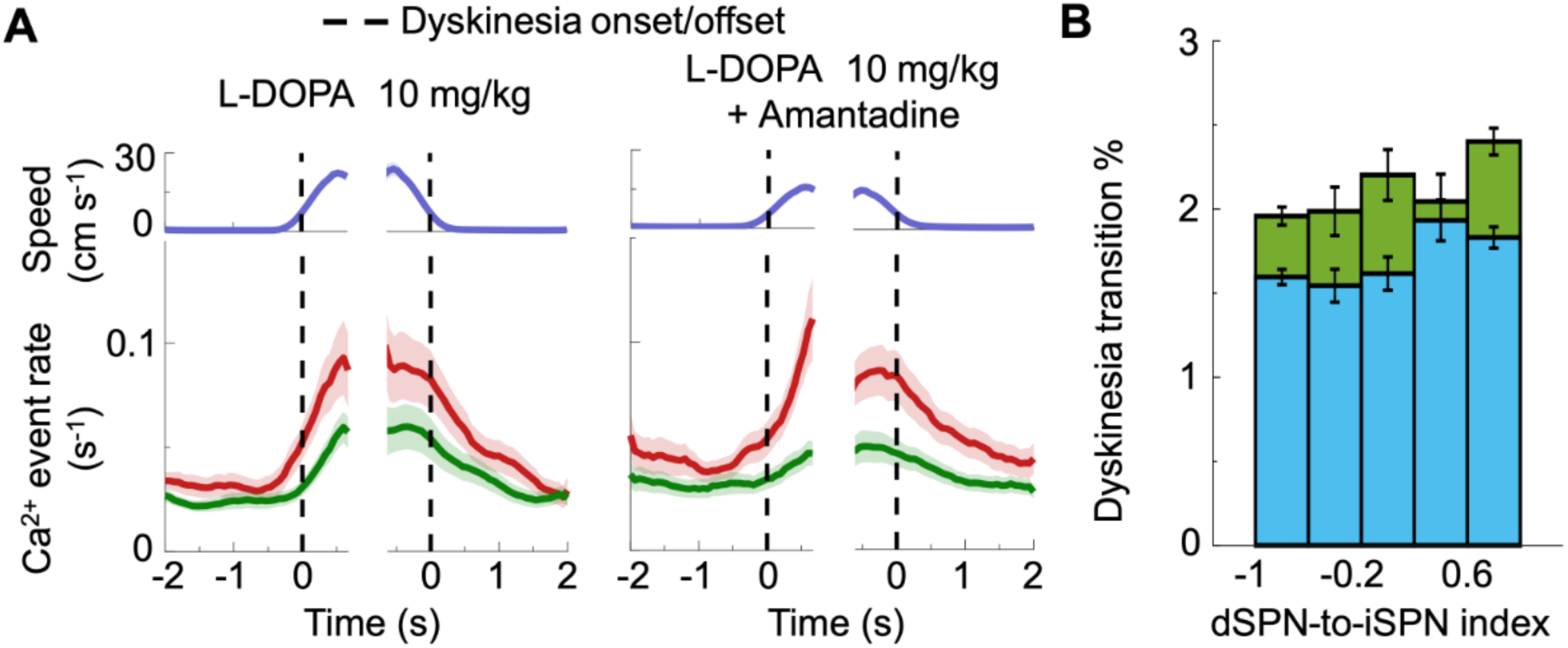
Dyskinesia-related SPN dynamics and dyskinesia initiation probabilities across conditions. (**A**) *Lower,* mean Ca²⁺ event rates of dSPNs (red) and iSPNs (green), aligned to dyskinesia onset and offset, shown for two treatment conditions. *Upper*, plots of mean forelimb speed. Shading: s.e.m. for n = 57 L-DOPA (10 mg/kg) blocks, and 55 L-DOPA (10 mg/kg) + amantadine blocks from 6 mice. **(B)** Bar plots showing the probability of transitioning from rest to dyskinesia relative to the dSPN-to-iSPN index of the rest period. The L-DOPA (10 mg/kg) condition had a significantly higher dyskinesia transition probability across all dSPN-to-iSPN index bins than the L-DOPA (10 mg/kg) + amantadine condition. Error bars: s.e.m. estimated using counting statistics for a binomial distribution (**Supp. Methods**).

**Supplemental Figure 9.**
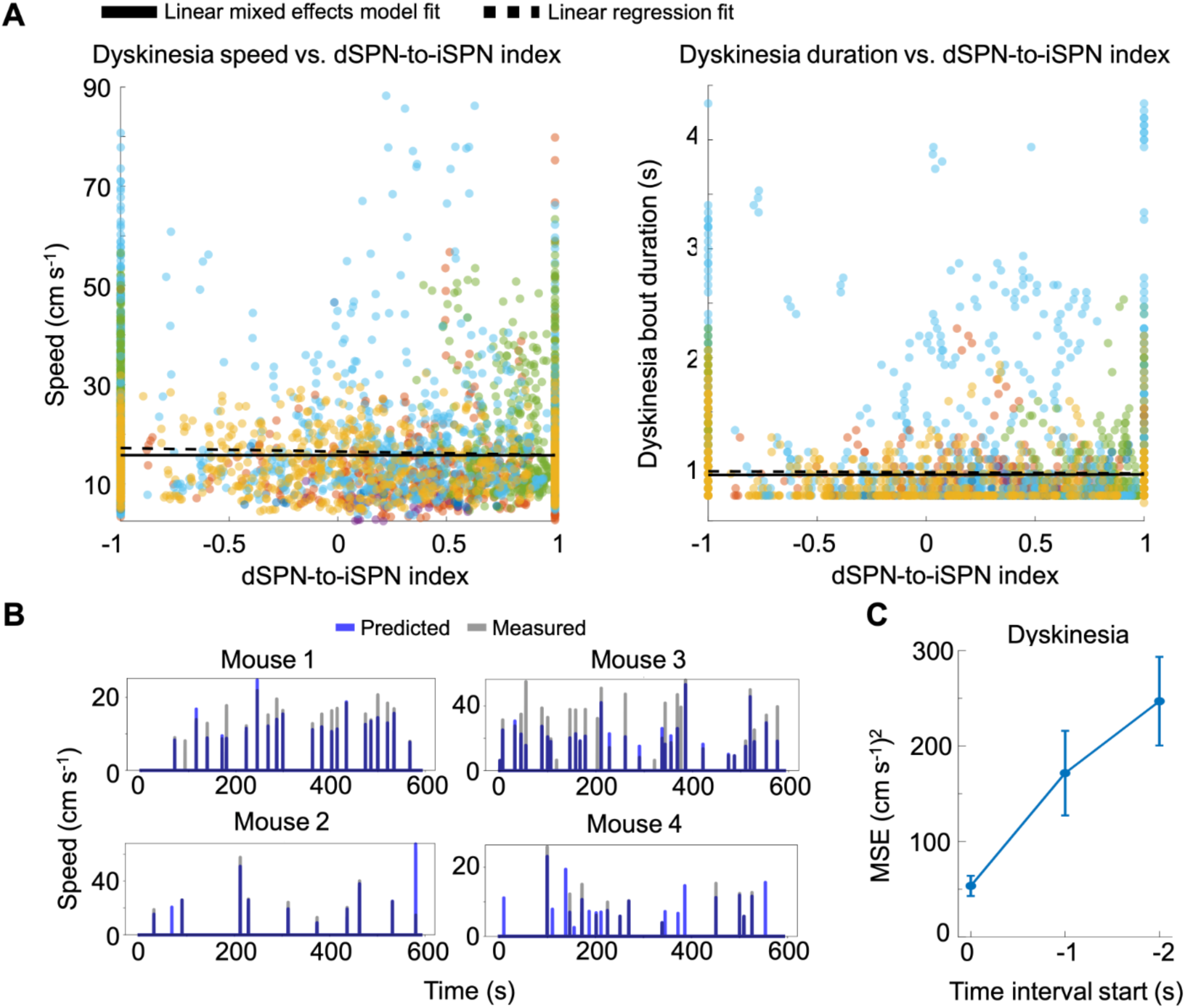
Neural ensemble activity patterns, but not the dSPN-to-iSPN index, predicts the kinematic attributes of upcoming dyskinesia bouts. (**A**) Scatter plots showing a lack of clear association between pre-dyskinesia dSPN-to-iSPN index values and either the speed (*left*) or duration (*right*) of upcoming dyskinetic movements. Each point denotes data from an individual bout of dyskinesia, with each color denoting an individual mouse. Data is pooled across 57 L-DOPA (10 mg/kg) blocks and 55 L-DOPA (10 mg/kg) + amantadine blocks. Solid black lines: Fits from a linear mixed-effects model, in which the systematic variations across mice were treated as random effects, and the variations across brain state and treatment conditions were treated as fixed effects. Dashed black lines: Fits from a least-squares regression applied to the pooled dataset without accounting for condition– or mouse-specific effects. For plots of dyskinesia forelimb speed *vs*. dSPN-to-iSPN index, the linear mixed-effects model yielded p = 0.94, and the pooled linear regression yielded R² = 0.003. For plots of dyskinesia duration *vs*. dSPN-to-iSPN index, the linear mixed-effects model yielded p = 0.77, and the pooled linear regression yielded a R² <0.001. **(B)** Example time series plots showing predicted (blue) and empirically measured (gray) forelimb speeds for dyskinetic movements, right after dyskinesia onset. Predicted speeds were based on the multivariate ridge regression model of **Figure 3D**. **(C)** Multivariate ridge regression performance results (**Figure 3D**, **Supp. Methods**) for different time intervals of 4-s-duration, shifted progressively further back in time from dyskinesia onset. The mean squared error (MSE) is plotted as a function of the start time of the 4-s-window, showing how the ability to predict the speed of dyskinetic forelimb movements based on neural activity patterns declines with increasing temporal separation from dyskinesia onset. Error bars: s.e.m. for n = 111 experimental blocks, comprising 57 L-DOPA (10 mg/kg) blocks and 54 L-DOPA (10 mg/kg) + amantadine blocks from 6 mice.

**Supplemental Figure 10.**
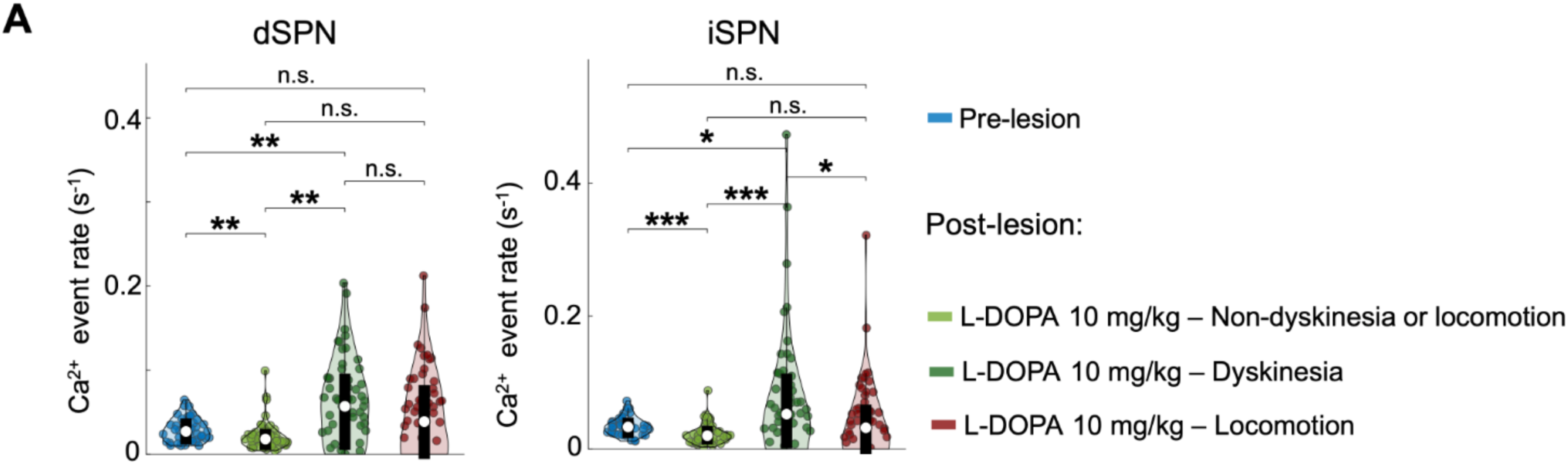
High-dose L-DOPA induces divergent activity patterns within the SPN population. (**A**) Violin plots showing the mean resting Ca²⁺ event rates of dSPNs and iSPNs in the pre-lesion state compared to three distinct SPN subpopulations during high-dose L-DOPA (10 mg/kg) treatment: those belonging to the dyskinesia ensemble, those belonging to the locomotion ensemble, and those belonging to neither (non-dyskinesia or locomotion). Outlier points > 3 s.d. from the mean not shown to avoid visual distortion of the plot. (*p < 0.05, **p < 0.01; two-sided rank sum tests corrected for multiple comparisons using a Benjamini–Hochberg procedure with a false-discovery rate of 0.05; n = 56 pre-lesion blocks and 57 L-DOPA (10 mg/kg) blocks from 6 mice).

**Supplemental Figure 11.**
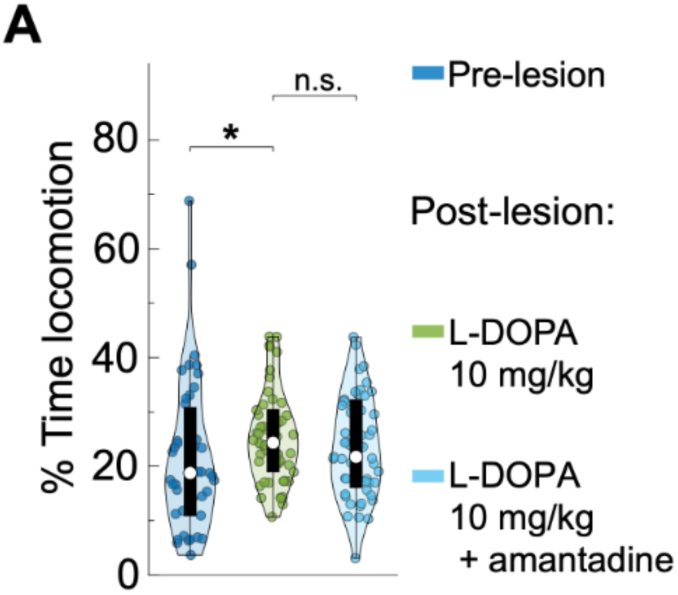
The percentage of time that dyskinetic mice spent locomoting was unaffected by amantadine treatment. (**A**) Violin plots of the percentage of time spent in locomotion for individual experimental blocks, across 3 different conditions. (**p < 0.01; two-sided rank sum tests corrected for multiple comparisons using a Benjamini–Hochberg procedure with a false-discovery rate of 0.05; n = 56 pre-lesion blocks, 57 L-DOPA (10 mg/kg) blocks, 56 L-DOPA (10 mg/kg) + amantadine blocks from 6 mice).

**Supplemental Figure 12.**
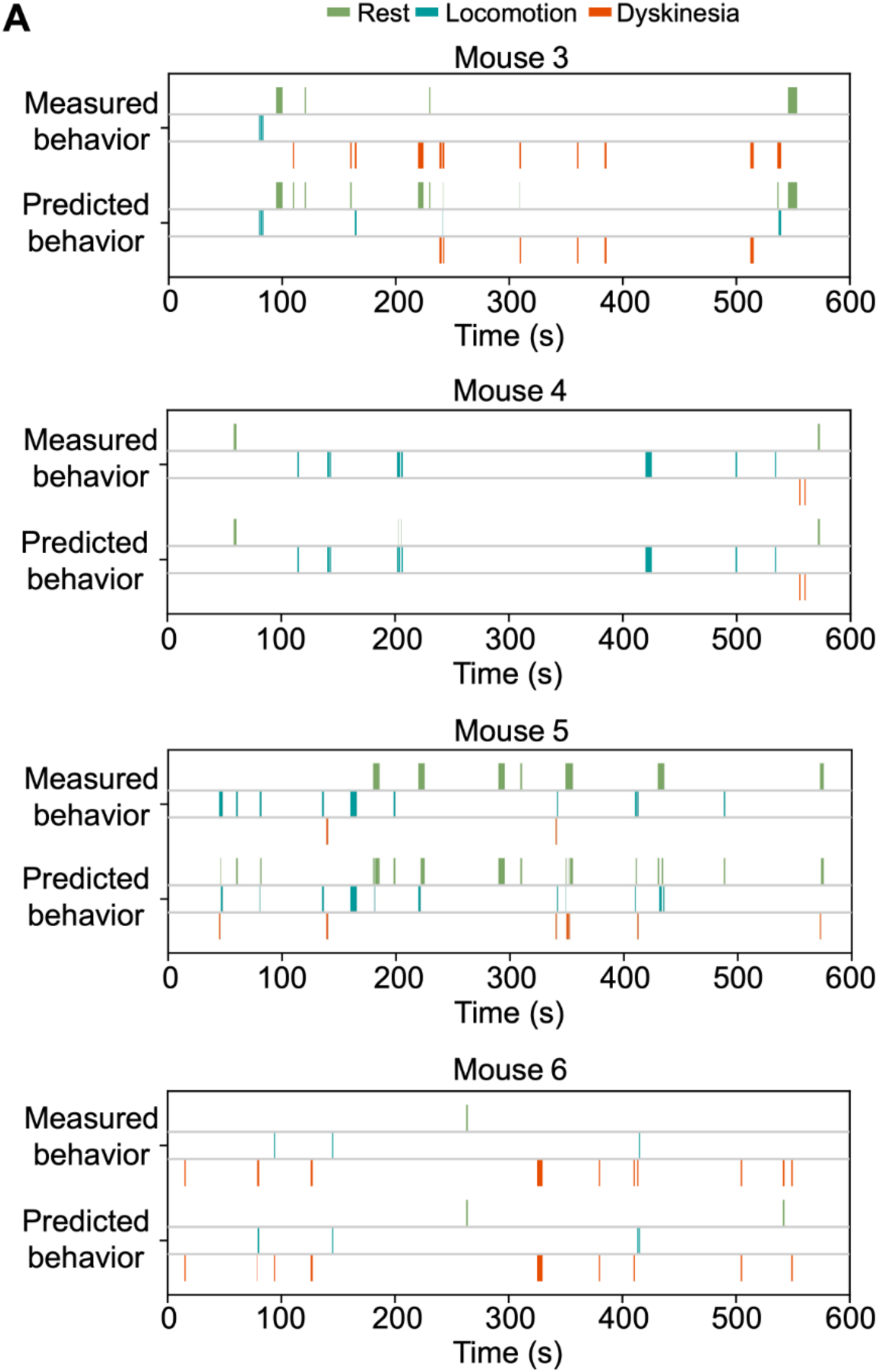
Distinct striatal activity patterns encode locomotion and dyskinesia. (**A**) Example time series in the same format as those of **Figure 4D**, showing example plots of empirically measured and SVM-predicted behavioral states for 4 additional mice.

**Supplemental Figure 13.**
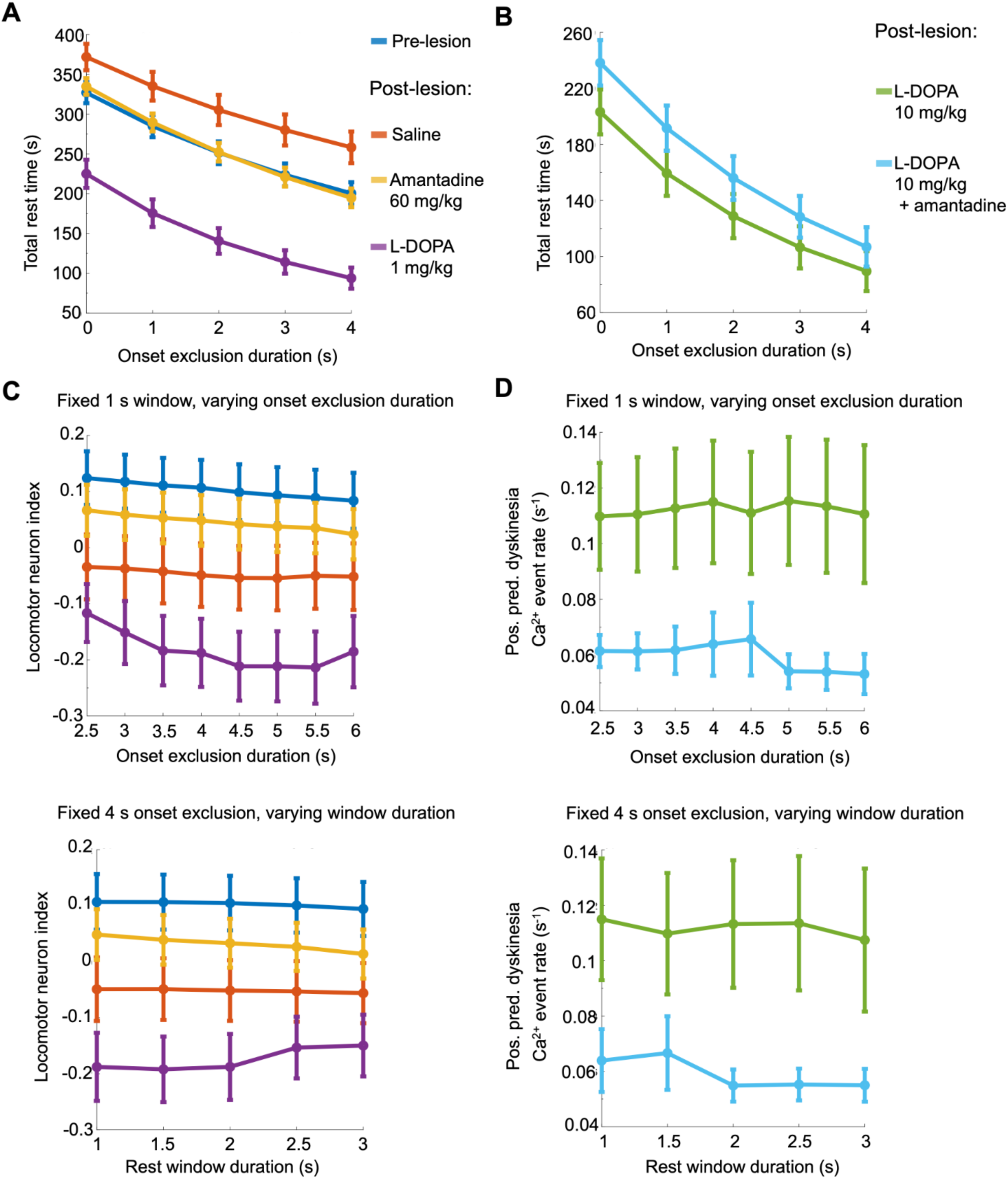
Rest time per block varies across conditions but remains substantial, and the main ensemble results are robust to changes in rest-period definition. (**A**) Mean rest time per block across hypokinetic conditions as a function of onset exclusion duration. Using the 1 s offset/onset requirement, mean retained rest remained substantial across conditions: pre-lesion, 285 s (48% retained); saline, 335 s (56% retained); amantadine, 290 s (48% retained); low-dose L-DOPA, 176 s (29% retained). With the more restrictive definition requiring an additional 4 s quiet period before movement onset, there was still substantial mean retained rest time: pre-lesion, 201 s (33% retained); saline, 258 s (43% retained); amantadine, 195 s (33% retained); low-dose L-DOPA, 94 s (16% retained). Error bars = s.e.m. (n = 50 pre-lesion blocks, 53 saline blocks, 53 amantadine blocks, and 37 L-DOPA (1 mg/kg) blocks from 5 mice). **(B)** Mean rest time per block across hyperkinetic conditions as a function of onset exclusion duration. Using the 1-s offset/onset requirement, mean retained rest was 160 s (27% retained) for high-dose L-DOPA and 192 s (32% retained) for high-dose L-DOPA + amantadine. With the more restrictive definition requiring an additional 4-s quiet period before movement onset, mean retained rest was 90 s (15% retained) for high-dose L-DOPA and 107 s (18% retained) for high-dose L-DOPA + amantadine. Error bars = s.e.m. (n = 57 L-DOPA (10 mg/kg) blocks and 54 L-DOPA (10 mg/kg) + amantadine blocks from 6 mice). **(C)** Mean resting locomotor neuron index per block as a function of onset exclusion duration (*top*) and rest window period duration (*bottom*) across hypokinetic conditions. Error bars = s.e.m. (n = 50 pre-lesion blocks, 53 saline blocks, 53 amantadine blocks, and 37 L-DOPA (1 mg/kg) blocks from 5 mice). **(D)** Mean resting positive-predictive dyskinesia neuron Ca²⁺ event rate per block as a function of onset exclusion duration (*top*) and rest window period duration (*bottom*) across hyperkinetic conditions. Error bars = s.e.m. (n = 57 L-DOPA (10 mg/kg) blocks and 54 L-DOPA (10 mg/kg) + amantadine blocks from 6 mice).

**Supplemental Figure 14.**
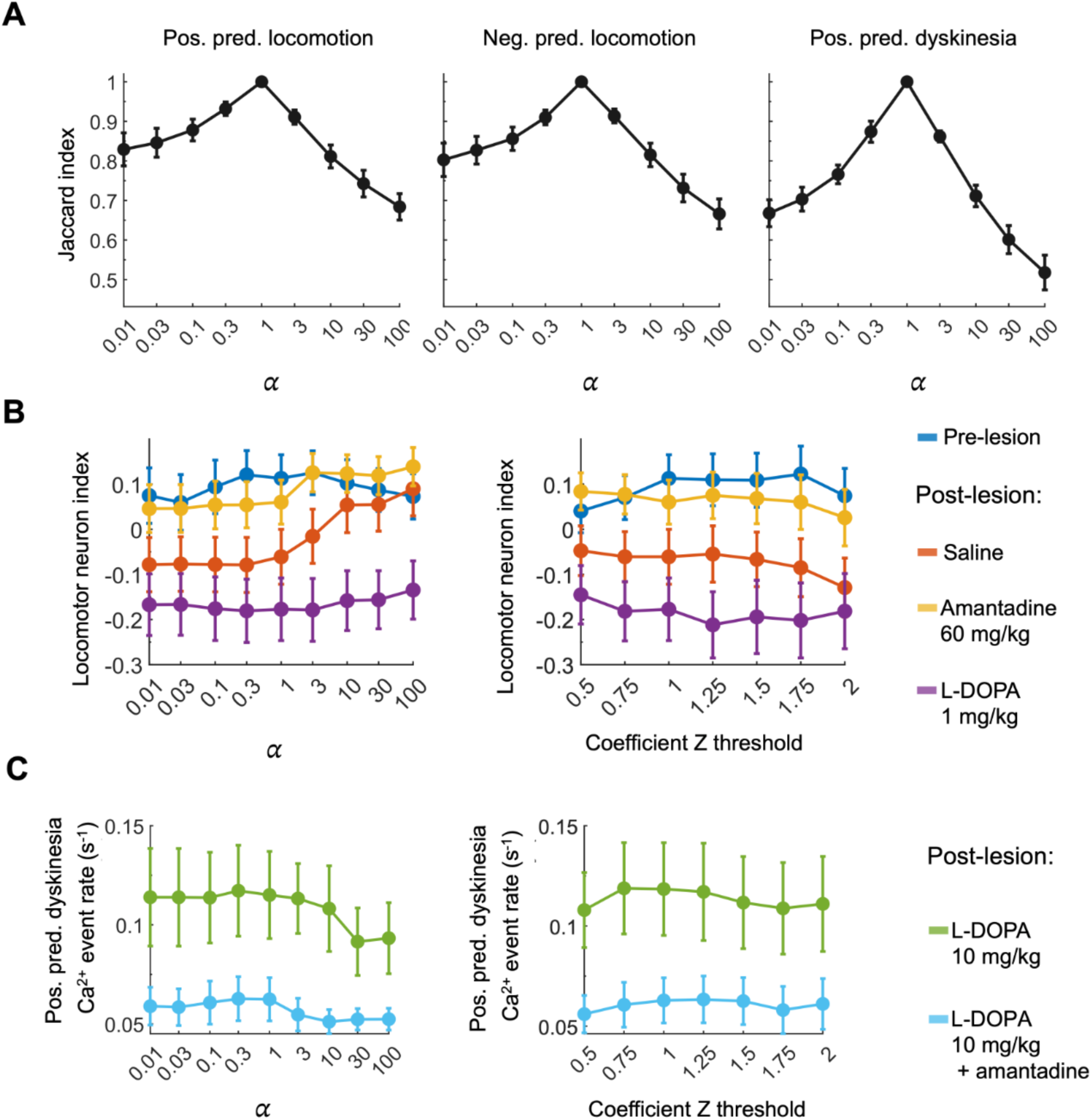
Ensemble identity and ensemble-level summary measure stability across changes in ridge-regression regularization coefficient threshold. (**A**) Jaccard index, relative to the default ridge-regression regularization parameter á = 1, across a logarithmic sweep of á values from 0.01 to 100, shown for positive-predictive locomotion ensembles, negative-predictive locomotion ensembles, and positive-predictive dyskinesia ensembles. Error bars = s.e.m. **(B)** Mean resting locomotor-neuron index per block as a function of á (*left*) and coefficient Z-threshold (*right*) across hypokinetic conditions. Error bars = s.e.m. (n = 50 pre-lesion blocks, 53 saline blocks, 53 amantadine blocks, and 37 L-DOPA (1 mg/kg) blocks from 5 mice). **(C)** Mean resting Ca2+ event rate of positive-predictive dyskinesia neurons per block as a function of á (*left*) and coefficient Z-threshold (*right*) across hyperkinetic conditions. Error bars = s.e.m. (n = 57 L-DOPA (10 mg/kg) blocks and 54 L-DOPA (10 mg/kg) + amantadine blocks from 6 mice).

## SUPPLEMENTAL METHODS

### EXPERIMENTAL MODEL AND STUDY PARTICIPANT DETAILS

#### Viral vectors

We obtained the plasmid, pGP-AAV-syn-jGCaMP7f-WPRE (#104488, Addgene)^32^, and subcloned the genetic sequence encoding jGCaMP7f from this plasmid into an AAV backbone with the CaMKII promoter, followed by the WPRE-hGH PolyA sequence, to create the plasmid pAAV-CaMKII-jGCaMP7f-WPRE.hGH. The Viral Tools facility at the HHMI Janelia Research Campus manufactured the virus AAV2/9-CaMKII-jGCaMP7f-WPRE.hGH, which we used at 5E12 GC/μL concentration to express jGCaMP7f in striatal neurons.

#### Pharmacological agents

In our study, we used the following pharmacological agents:

● 6-Hydroxydopamine hydrobromide (6-OHDA): Obtained from Fisher Scientific (50-194-8364; Purity: ≥99.9%). Solvent: Dissolved in 0.9% saline with 0.02% L-ascorbic acid (A92902, Sigma-Aldrich).
● Levodopa (L-DOPA): Obtained from Sigma-Aldrich (PHR1271; Purity: ≥99.7%). Solvent: Dissolved in 0.9% saline.
● Benserazide hydrochloride: Obtained from Sigma-Aldrich (B7283; Purity: ≥98%). Solvent: Dissolved in 0.9% saline.
● Amantadine hydrochloride: Obtained from Sigma-Aldrich (A1260; Purity: ≥98%). Solvent: Dissolved in 0.9% saline.
● Desipramine hydrochloride: Obtained from Fisher Scientific (30-675-0; Purity: >99%). Solvent: Dissolved in 0.9% saline.

The 0.9% saline solution (Hospira) contained 154 mEq/L of sodium (Na⁺) and 154 mEq/L of chloride (Cl⁻), with a calculated osmolarity of 308 mOsmol/L. To ensure their stability and efficacy, we prepared all solutions fresh on the day of the experiment.

#### Mice

The Stanford Administrative Panel on Laboratory Animal Care (APLAC) approved all experiments. To facilitate concurrent recordings and the differentiation of direct pathway spiny projection neurons (dSPNs) and indirect pathway spiny projection neurons (iSPNs), we used a mouse line expressing the red fluorophore tdTomato specifically in dSPNs. We first crossed homozygous Drd1a^cre^ mice^30^ with homozygous Ai14 reporter mice (Gt(ROSA)26Sor^tm14(CAG-tdTomato)Hze^; Allen Brain Institute)^31^, which express the red fluorophore tdTomato in a Cre-dependent manner, to generate double heterozygous offspring. We intercrossed the double heterozygous mice to produce homozygous Ai14 / homozygous Drd1a^cre^ mice, which then served as breeders for all mice used in the experiments. We previously used the same type of double transgenic mice in an earlier study.^8^ Here, our data came from 3 female and 3 male mice, aged 16–36 wks at the start of behavioral experiments. One animal was excluded from hypokinetic condition analyses because it had inadvertent L-DOPA exposure before the saline (control) recording session, which violated the intended session order (see **Drug treatments**). However, as it successfully developed L-DOPA-induced dyskinesias, this animal was included in the hyperkinetic condition analyses. Thus, N = 5 mice for hypokinetic condition comparisons and N = 6 for hyperkinetic condition comparisons. We housed mice in 12-h light/dark cycle conditions. We group housed mice (2–5 per cage) before surgery and housed them individually after surgery.

### EXPERIMENTAL METHODS

#### Surgeries

To enable repeated imaging of striatal neurons and to model a parkinsonian state, we performed the surgeries described below. For all surgeries, we anesthetized mice with isoflurane (1-4% in O_2_) and positioned them within our stereotaxic apparatus using ear bars (Model 963, David Kopf instruments).

To express a genetically encoded Ca^2+^ indicator (jGCaMP7f)^32^ in SPNs of the dorsolateral striatum (DLS), we performed a stereotaxic injection of 500 nL AAV2/9-CaMKII-jGCaMP7f (5E12 GC/μL) into the right DLS (anteroposterior (AP): +0.5 mm from bregma; mediolateral (ML): +2.3 mm from bregma; dorsoventral (DV): –2.3 mm).^19^ We performed injections with an UMP3 Ultramicropump (UMP3-3, World Precision Instruments) and NanoFil syringe with a 33-gauge beveled tip (NF33BV, World Precision Instruments). Virus injections were performed at a rate of 200 nL/min. After injections, we sutured the scalp and allowed mice to recover for 1 week.

To allow *in vivo* imaging of SPN activity in the DLS, we removed the scalp and used a 1.4-mm-diameter drill bit to create a craniotomy over the right DLS. Using a 27-gauge needle attached to wall-mounted vacuum tubing, we aspirated the overlying brain tissue and implanted an optical guide tube into the craniotomy site with the end of the tube at a depth of –2.0 mm DV. We created the optical guide tube by gluing a ∼1.1 mm diameter disc of 0.1 mm Schott glass (custom fabricated, TLC International) to the tip of a 3.8-mm-long, 18-gauge stainless steel tube (custom fabricated, Ziggy’s Tubes and Wires). After the guide tube was in place, we placed a 1-mm-diameter, 4-mm-long gradient index (GRIN) lens (NA 0.5, Pitch 0.5, 1050-004595, Inscopix) into the tube, and we attached a steel headbar to the skull to allow head-fixation. We affixed the full assembly (head bar, optical guide tube with GRIN lens) to the skull by applying first an ultraviolet-light-curing adhesive (Loctite #4305, IDH:303389, R.S. Hughes) and then dental cement (Stoelting Dental Cement, #10000786, Fisher Scientific). We left an open area over the right posterior aspect of the skull (an approximately circular area with a 2-mm-diameter centered on AP: –3.0 mm; ML: +1.3 mm) to facilitate the subsequent injection of 6-OHDA to the substantia nigra pars compacta (SNc).

Mice recovered for 3 weeks after this implantation surgery before starting imaging and behavioral experiments. After completing pre-lesion behavioral and neural recordings, we induced a hemiparkinsonian state by performing a unilateral 6-OHDA lesion to selectively ablate dopaminergic neurons in the SNc. To prevent lesioning of non-dopamine monoamine neurons, we first injected mice with desipramine (25 mg/kg; i.p.; Fisher Scientific) 20 min before 6-OHDA injections. We then stereotaxically injected 6-OHDA (2 μg/μL, dissolved in 0.02% L-ascorbic acid in 0.9% saline; 6-OHDA from Fisher Scientific, L-ascorbic acid from Sigma-Aldrich) in the right SNc (AP: –3.0 mm; ML: +1.3 mm) at 3 depths below the dura (–4.2, –4.0, and –3.8 mm DV; 1.3 μL of 6-OHDA solution at each depth).^8^ Mice recovered for at least 2 weeks before starting the next round of experiments (post-lesion behavior and neural recordings).

#### Behavioral recordings

We head-fixed mice and placed them on a custom 3D-printed running wheel (a 4.3-cm-diameter, 10-cm-long cylinder) equipped with a rotary encoder that provided wheel position information. We positioned a camera (DMK37BUX273, The Imaging Source) and infrared illuminator (AI4, Tendelux) in front of the mice to record behavior, such as forelimb movements. A custom hardware and software setup (National Instruments myRIO, LabVIEW 2019) time-locked the neural recordings with the behavioral readouts from the rotary encoder and camera. For each image frame recorded by the two-photon microscope (30 fps), one measurement was recorded from the rotary encoder (making it a 30-Hz-recording) and 3 frames were recorded by the camera, making it a 90-fps-recording. See ‘Two-photon Ca^2+^ imaging’ below for more details.

#### Two-photon Ca^2+^ imaging

Once mice were implanted with the optical guide tube and GRIN lens in the dorsolateral striatum (see ‘Surgeries’), we acquired Ca^2+^ imaging data from 3 different axial planes (at least 20 μm apart in depth) in the brains of head-fixed mice free to walk or run in place on a wheel. During each imaging session, we imaged the 3 planes sequentially (plane 1, then plane 2, then plane 3) and performed this sequence twice, yielding two recordings per plane. For each session, we positioned head-fixed mice such that the GRIN lens was placed under the objective lens (NA 0.8, CFI75 LWD 16X W, Nikon) of a custom-built two-photon microscope.

We performed two-photon imaging using ultrashort-pulsed laser illumination (940 nm wavelength, to excite both GCaMP and tdTomato) emitted by a wavelength-tunable laser (MKS Spectra-Physics, Insight X3, with dispersion compensation). We used two-stage control of the laser illumination power, with an achromatic quarter-wave plate (Thorlabs, AQWP10M-980, 690–1200 nm) and a polarization-dependent beam splitter to provide coarse attenuation, followed by an electro-optical modulator (EOM; Conoptics) to provide fine control and fast blanking. Depending on the specific setup, either a Conoptics EOM 350-80-LA-02 with a 302RM driver or a Conoptics AP214 EOM was used for high-speed modulation and rapid laser blanking during galvanometer turnaround.

Laser beam scanning was performed by two galvanometric mirrors, including a resonant scanner (8 kHz) for the fast-scanning axis (Novanta Photonics, formerly Cambridge Technology, CRS8K 6215H 6SD11015 SYS 2X). Fluorescence emissions were directed onto gated photomultiplier tubes (PMTs; Hamamatsu, H11706P-40 and H10770PA-40 for green and red channels, respectively). The microscope objective lens was a Nikon 16×, NA = 0.8, with a 3 mm working distance (WD). A custom overhead collimated laser diode (532 nm wavelength, Thorlabs, CPS532) was used to check the proper mechanical alignment of the mouse by monitoring the back reflections of the diode laser beam from the sample GRIN. The mouse stood on a custom platform equipped with two goniometers to maintain a fixed orientation. We operated the microscope using a customized version of ScanImage software (version 5.6.1, MBF Biosciences, formerly Vidrio Technologies), which enabled real-time image acquisition (∼30 fps, 512 × 512 pixels), hardware control, axial focusing, z-stack collection, and custom metadata saving. Customizations included X-Y sample translation, provided by a motorized Sutter MP-285 stage, and fine axial (Z) control performed by a piezoelectric stage (PI, Q-545.240) with an E-873.1AT controller.

Synchronization of the two-photon microscope (∼30 Hz frame acquisition rate) and the faster behavioral camera was achieved using a custom solution based on a National Instruments MyRIO FPGA. The MyRIO detected rising edges of the two-photon frame clock, measuring the interval between consecutive edges to determine the nominal period (T). It generated a train of short (∼100 µs) electronic pulses, subdividing T into m equal intervals (typically m=3), and transmitted these pulses through digital I/O lines as frame triggers for the behavioral camera. The clock upsampling yielded a simple relationship between the number of two-photon image frames (n) and the total frames acquired by the behavioral camera ((n-1)×m). Each frame from the behavioral camera was time-locked to the two-photon image sequence, eliminating a need for offline resampling or temporal alignment. Real-time communication between the MyRIO and the microscope software was managed via built-in LabVIEW interfaces and FIFO communication. Trigger signals and timestamps were logged in tab-separated values files provided by a PC host computer operating the MyRIO.

#### Drug treatments

To assess the effects of different pharmacological interventions on motor behavior and neuronal activity in the parkinsonian state, 2–8 wks after the 6-OHDA lesion we began post-lesion experimental sessions. 20 min before each set of ‘post-lesion’ session, we injected mice with either: 0.9% saline (post-lesion control), amantadine (60 mg/kg), L-DOPA (1 mg/kg) + benserazide (15 mg/kg), L-DOPA (10 mg/kg) + benserazide (15 mg/kg), or L-DOPA (10 mg/kg) + benserazide (15 mg/kg) + amantadine (60 mg/kg) (amantadine, L-DOPA, and benserazide all from Sigma-Aldrich). In all sessions that we administered L-DOPA, we co-administered benserazide to prevent peripheral conversion of L-DOPA to dopamine. We selected doses based on prior studies.^8,25^ We dissolved all compounds in 0.9% saline and delivered them intraperitoneally (i.p.) with a 30 mL/kg injection volume. We performed drug treatment sessions 1–10 d apart. We found that the ordering of drug treatments was very important, as injections of L-DOPA would prime the mice for dyskinesias, and mice could then have dyskinesias even without L-DOPA injections. Therefore, saline-only sessions were always done first, then amantadine only sessions, and then L-DOPA (1 mg/kg) sessions. Finally, we alternated L-DOPA (10 mg/kg) and L-DOPA (10 mg/kg) + amantadine sessions, with the first session of L-DOPA (10 mg/kg) always occurring before the first L-DOPA (10 mg/kg) + amantadine session (**Figure 1E**). One mouse inadvertently received L-DOPA prior to its saline-only session and was therefore excluded from hypokinetic comparisons (see **Mice**). During each session, there were 6 blocks of two-photon imaging and behavioral recording, each 10 min in duration, corresponding to two sequential passes through 3 different axial planes in the DLS.

#### Immunohistochemistry

After the completion of imaging experiments, we euthanized and intracardially perfused the mice with phosphate-buffered saline (PBS) (Gibco) and then a 4% solution of paraformaldehyde (Electron Microscopy Sciences) in PBS. We sliced the fixed brain tissue using a vibratome (VT1000s, Leica) to obtain 80-μm-thick coronal sections. We immunostained the tissue sections with polyclonal antibodies against tyrosine hydroxylase (TH 1:500, Aves Labs) and applied the secondary antibody Alexa Fluor 647 goat anti-chicken IgG(H+L) (A-21449, Thermo Fisher). Sections were also stained with the nuclear counterstain DAPI (D1306, Invitrogen). We visualized immunofluorescence in our sections using a slide-scanning microscope (Axio Imager Z1, Zeiss) and confirmed that all mice included for analyses had a complete unilateral loss of dopaminergic signal in the striatum and substantia nigra pars compacta (SNc) (**Figure S1A**).

### QUANTIFICATION AND STATISTICAL ANALYSIS

#### Behavioral analyses

We performed automatic limb position tracking from the behavioral video recordings using DeepLabCut (version 2.0).^40^ We classified locomotion periods as intervals when the speed of the wheel was > 0.5 cm/s. We classified dyskinesia periods as intervals when the left forelimb, contralateral to the 6-OHDA lesion in all mice, was moving at an average speed > 2.5 cm/s (with at least 3 frames during the period reaching > 30 cm/s) and when there was no sign of locomotion on the running wheel (**Figures 1C,D**). For behavioral analyses, we defined the following types of periods: **pre-locomotion**, a 2-s-period preceding locomotion onset; **pre-dyskinesia**, a 2-s-period preceding dyskinesia onset; and **rest**, periods of inactivity at least 1 s after locomotion or dyskinesia offset and at least 1 s before locomotion or dyskinesia onset. To analyze the resting activity of locomotion and dyskinesia predictive neurons, we excluded rest periods that occurred within 4 s of locomotion or dyskinesia onset. This ensured that data intervals used in multivariate regression analyses for the identification of neural activity patterns predictive of upcoming locomotion or dyskinesia were not reused in the subsequent analyses of the resting activity of these same neurons. Condition-specific retained resting time and robustness to alternative resting-window definitions for the principal ensemble analyses are summarized in **Figure S13**.

Our dyskinesia analysis focused on forelimb dyskinesias as these were robustly seen in all mice. Orolingual dyskinesias were clearly identified only in 2 of 6 mice; in the remaining animals, facial movements could not be reliably distinguished from baseline, precluding cohort-level quantification.

#### Analysis of Ca^2+^ video datasets

We pre-processed Ca^2+^ videos by temporally downsampling each movie twofold and then corrected computationally for rigid and non-rigid lateral motion of the brain using the NoRMCorre image registration routine.^49^ To obtain activity traces of individual neurons’ relative changes in [Ca^2+^]-related fluorescence (Δ*F*(*t*)/*F*) from the motion-corrected Ca^2+^ videos, we used the cell extraction routine EXTRACT.^50,51^ From each neuron’s extracted Ca^2+^ activity trace, we identified Ca^2+^ transient events via detection of fluorescence peaks that were ≥ 2 s.d. above the cell’s mean baseline fluorescence level and lasted for ≥ 0.2 s. We determined the time of each Ca^2+^ event as the temporal midpoint between the time of the Ca^2+^ event’s fluorescence peak and the most recent preceding trough in fluorescence. To classify neurons found by EXTRACT as either dSPNs or iSPNs, we first used the software Cellpose to determine the outlines of all cells expressing tdTomato.^52,53^ Each neuron determined by EXTRACT to have a spatial profile within the outline of a tdTomato-expressing-cell was classified as a dSPN. We classified all other neurons found by EXTRACT as iSPNs.

#### Identification of behavior-predictive neurons

To identify striatal neurons with activation patterns predictive of either locomotion or dyskinesias, we performed linear regression analyses to identify SPNs the activity of which predicted the mouse’s upcoming movement speed.

To do this, we built predictive models that linked neural activity to the speed of either locomotion or dyskinetic forelimb movements (**Figures 2C, 3D**, **S3B**, **S6B**). For each behavioral block, we first identified the onset time of each locomotion or dyskinesia bout and measured the initial movement speed over the first 0.66 s following movement onset. This value served as the behavioral response variable. We also included 10 periods of inactivity (*i.e.*, when neither locomotion nor dyskinesia occurred) as additional response variables in the dataset.

For each behavioral onset, we used SPN activity from the interval [–4 s, 0 s] just prior to movement onset to create a predictor matrix. Each row in this matrix corresponded to a single movement or rest period. Each column of the matrix contained the mean Ca^2+^ event rates of an individual neuron across the set of pre-movement periods. This approach allowed us to examine how neural activity prior to movement related to the upcoming behavioral outcome.

To account for differences in the scale and distribution of neural activity, we z-scored all predictor variables and applied a log-transformation to the response variable to mitigate skewness. We then used a ridge regression model (**cuml.linear_model.Ridge** module, RAPIDS AI framework) to predict the speed of locomotion or dyskinesia forelimb movements based on the neural activity patterns. We chose ridge regression for its capability to handle multicollinearity between neurons while applying *L*_2_ regularization. We trained the regression model separately for each dataset using a regularization parameter (alpha) of 1.0, and we evaluated its performance based on its mean squared error (MSE) and the coefficient of determination (R²) on the training dataset. The sensitivity of ensemble assignment and the main downstream results to the choice of regularization parameter is shown in **Figure S14.** To quantify the stability of ensemble membership across regularization parameters, we computed the Jaccard index between the ensemble identified at each tested alpha value and the ensemble identified using the default alpha = 1, where the Jaccard index is the size of the intersection divided by the size of the union of the two ensembles.

To identify behavior-predictive neurons, we examined the coefficients determined for each neuron in the ridge regression models. We classified neurons with strong positive regression coefficients (z-score > +1 above those of the mean across all neurons) in the locomotion models as positive-predictive locomotor neurons and those with strong negative coefficients (z-score < –1) as negative-predictive locomotor neurons. Similarly, we classified neurons with strong positive coefficients (z-score > +1) in the dyskinesia models as positive-predictive dyskinesia neurons. We used this threshold as a practical compromise to identify relatively strong coefficient weights without making the resulting ensembles overly sparse or overly inclusive. The sensitivity of the main downstream treatment-direction effects to this threshold choice is shown in **Figure S14**. We validated the behavioral associations of these neuron classes by comparing the speeds of locomotion or forelimb movement in the interval [0 s, 0.66 s] prior to Ca²⁺ event onset to those in the interval [3.33 s, 4 s] after onset. We separately performed regression analyses for identifying positive-predictive locomotion and dyskinesia neurons, implying that, in principle, individual neurons could be classified as both, even though few were (**Figure 4C**).

Our main analyses focused on the [–4 s, 0 s] interval just prior to movement. However, to test the extent to which neural signals predictive of movement arose before movement onset, we conducted ancillary analyses in which we progressively shifted the predictor window backwards in time from movement onset (**Figures S5C**, **S9C**). As expected, predictive performances declined as the prediction window moved further in time from the moment of movement onset.

We constructed regression models separately for each mouse, drug condition, and optical plane. For each of these datasets, we included the two recording blocks performed at each optical plane, and we tracked individual neurons across blocks whenever possible based on spatial alignment. Not all neurons were present in every block, which could in principle bias the determinations of regression coefficients. However, the z-scored regression coefficients for each neuron did not meaningfully correlate with the number of blocks in which the neuron was detected, suggesting that coefficient strength was not simply driven by the number of blocks in which we recorded a cell’s activity. This approach allowed us to include more behavioral bouts per neuron, with the aim of improving the accuracy with which the regression coefficients could be determined.

#### Measures of relative SPN activity

To quantify the relative levels of dSPN and iSPN activity, we computed a dSPN-to-iSPN index, defined as the difference between the mean Ca^2+^ event rates for dSPNs (*Rate_dSPN_*) and iSPNs ( *Rate_iSPN_*), divided by their sum: (*Rate_dSPN_* – *Rate_iSPN_*)/(*Rate_dSPN_* + *Rate_iSPN_*). However, we noted a sampling bias in our dataset wherein more iSPNs than dSPNs had detectable Ca^2+^ activity (78 ± 11% of neurons with detectable Ca^2+^ activity were iSPNs). To ensure our dSPN-to-iSPN index was robustly calculated and not confounded by this unequal cell count, we employed a 1:1 subsampling bootstrap procedure. For each bout, we identified the total number of available dSPNs and iSPNs and found the minimum of the two (*n_min_*). If *n_min_* > 0, we performed 1,000 bootstrap iterations, randomly selecting *n_min_* dSPNs and *n_min_* iSPNs from their respective total pools and calculated a dSPN-to-iSPN index. We then averaged over all 1,000 iterations for the final, bias-corrected dSPN-to-iSPN index value used for analysis. To assess the numerical stability of this bootstrap equalization procedure, we quantified the standard deviation of the dSPN-to-iSPN index across the 1,000 bootstrap resamples for the analyses underlying Figures 2A and 3B. Across conditions, the mean bootstrap standard deviation ranged from 0.09 to 0.22, indicating that the estimated index was not dominated by large resampling instability. We did not emphasize the coefficient of variation because this metric becomes unstable when the mean index is near zero.

To quantify the relative activity levels of positive– and negative– predictive locomotion neurons, we calculated a locomotor neuron index, defined as the difference between the mean Ca^2+^ event rates for positive– (*Rate_positive_Locomotion_*) and negative– predictive locomotion neurons (*Raten_egative_Locomotion_*), divided by their sum:

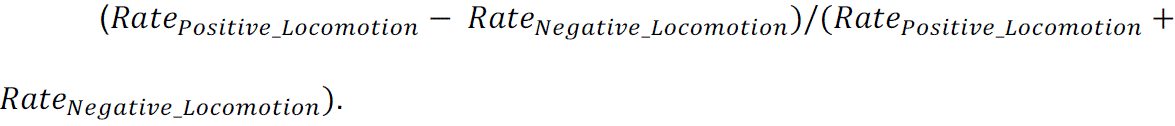

#### Linear mixed-effects model

To analyze whether the resting state activity levels of specific neurons were predictive of the extent of mouse movement, we applied linear mixed-effects models across multiple brain states, treatment conditions and individual mice (**Figures 2I**, **3L, S7B**). In these models, we included the neural activity metric of interest (*e.g*., dSPN-to-iSPN index, locomotor neuron index, or positive-predictive dyskinesia neuron mean Ca^2+^ event rate) as a continuous fixed effect, the experimental condition as a categorical fixed effect, and mouse identity as a random effect that accounted for inter-mouse variability. For locomotion and dyskinesia behaviors, we specified the models as:

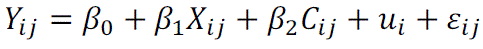

where:

*Y_ij_* = Percentage of time spent in locomotion or dyskinesia for mouse *i* in block *j*

*X_ij_* = Neural activity metric (*e.g.*, dSPN-to-iSPN index) for mouse *i* in block *j*

*C_ij_* = Condition (categorical fixed effect)

*u_i_* = Random intercept for mouse *i*

*ε_ij_* = Residual error

We fit the models using MATLAB’s (R2023a, MathWorks) linear mixed-effects modeling framework (**fitlme**) and optimized the parameter estimates by maximum likelihood estimation. This additive model was used as the primary analysis to test whether a general association was present between neural activity and behavior across treatment conditions. To further characterize the contribution of between-animal variability, we examined the intraclass correlation coefficient (ICC), defined here as the fraction of total variance attributable to differences between mice. The ICC values were approximately 0.39-0.43 for the hypokinetic models and 0.69-0.70 for the hyperkinetic models, indicating substantial between-animal variability, especially in the hyperkinetic analyses. Because these models were fit across imaging blocks from only 5-6 animals, the mouse random effect was estimated from a modest number of levels, and the mixed-effects results should therefore be interpreted with caution.

In addition, we used further linear mixed-effects analyses (**Figures S5A, S9A**) to determine, at the level of individual movement bouts, whether the dSPN-to-iSPN index during the pre-movement period was predictive of the kinematic attributes of the upcoming locomotion or dyskinesia bouts. Specifically, we tested whether the dSPN-to-iSPN index in the seconds before movement onset correlated with the speed or duration of the ensuing movement bout. These linear mixed-effects models were constructed similarly as those described above, with the pre-movement dSPN-to-iSPN index used as the neural activity metric (*X*) and the upcoming locomotion or dyskinesia speed or duration as the outcome variable (*Y*).

#### Behavioral state decoding using support vector machines (SVMs)

To determine whether distinct behavioral states—rest, locomotion, and dyskinesia—are separably represented in the dorsolateral striatum (DLS), we trained support vector machine (SVM) decoders to classify behavioral states based on neural activity (**cuml.svm.SVC** module, RAPIDS AI framework). This process involved extracting neural activity features, labeling behavioral states, training the model, and evaluating its performance.

To link neural activity with specific behaviors, we first segmented each experimental block into overlapping 1-s-long time windows, with the center of each window spaced 0.33 s apart. We assigned a behavioral label (rest, locomotion, or dyskinesia) to each window based on the movement measurements during that time period. A time window was classified as locomotion if the mouse exhibited continuous locomotion for the entire 1 s (without simultaneously detected dyskinesias). A time window was classified as dyskinesia if the mouse exhibited dyskinetic movements for the entire 1 s (see **Behavior** for dyskinesia identification details). To exclude periods with potential transitions between behavioral states, a window was classified as a rest period if neither locomotion nor dyskinesia occurred during the 1 s window itself or within 5 s buffer periods before and afterward. Time windows that did not meet any of the above criteria were not included in the datasets used to train or test the SVM decoders.

For each neuron, we represented its activity within a given time window using the cell’s mean Ca^2+^ event rate. This yielded a feature vector, in which each element denoted an individual neuron’s net activity within the time window. For each experimental block, 24 randomly selected 5-s-long periods were designated as the test set (total of 120 s, 20% of the block), and the model was trained on all remaining data (480 s, 80% of the block). This approach was chosen over pure random sampling to minimize information leakage from temporal correlations of the Ca^2+^ indicator. To balance the class distributions, we applied random undersampling to the training dataset. To assess classification performance while accounting for variability across the train-test splits, we repeated model training and testing 100 times per imaging plane and mouse. In each iteration, we trained a support vector classifier on the undersampled training data and evaluated its performance on the held-out test dataset. We then averaged the results across all 100 train-test splits and quantified classification accuracy. To establish chance-level performance, we trained a parallel decoder on shuffled training set labels in each of the 100 iterations. To illustrate classification performance, we generated example decoded time series and plotted the empirically determined mouse behavioral periods alongside those predicted by the SVM for individual experimental blocks (**Figures 4D, S12A**). To determine if decoder performance was dependent on the size of the recorded neural population, we analyzed the decoders’ accuracy versus the total number of recorded neurons per block. We then pooled this data from all mice and blocks and grouped it into six bins based on the number of neurons. The mean accuracy and standard error of the mean were calculated for each bin to visualize the relationship between recorded neuron number and decoding performance (**Figure 4E**).

#### Statistical analysis

For two-sample comparisons of a single variable, we used a non-parametric test, either the Wilcoxon rank sum or the Wilcoxon signed-rank test (MATLAB functions, **ranksum** and **signrank**). All tests were two-tailed unless stated otherwise. Where applicable, we corrected for multiple comparisons using a Benjamini–Hochberg procedure with a false-discovery rate of 0.05. To determine statistical significance in the linear mixed-effects models, we used the p-value estimated by the model. Detailed statistical results are provided in Supplemental Table 1. For Wilcoxon analyses, the table reports Hodges–Lehmann shift estimates with corresponding nonparametric confidence intervals; for one-sided Wilcoxon tests, one-sided confidence intervals are reported in the direction of the prespecified alternative hypothesis. For linear mixed-effects models, the table reports the fixed-effect coefficient (beta) and its 95% confidence interval. For ordinary least-squares regression analyses, the table reports the slope and its 95% confidence interval.

For our violin plots, the shape of the violin illustrates the kernel density estimate (KDE), providing a smoothed representation of the data distribution. This KDE was calculated using MATLAB’s default **ksdensity** function, which uses a Gaussian kernel to estimate the probability density function of the data. See also the caption of **Figure 1** for further details about the plot format.

For transition probability barplots (**Figures S3C, S8B**), each bin’s transition probability was calculated as 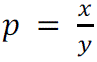 where *x* is the number of transition trials and *y* is the total trials in that bin. The probabilities are plotted as percents (*p* × 100). Error bars indicate the s.e.m. of this proportion, determined as 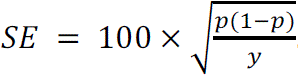.

#### Software

We used ScanImage software (version 5.6.1, MBF Biosciences, formerly Vidrio Technologies) and custom Labview 2019 software (National Instruments) to record two-photon Ca^2+^ videos. We used NoRMCorre for image registration of the Ca^2+^ video frames and EXTRACT for identifying neurons in these videos and determining their fluorescence activity traces.^49–51^ We used Cellpose to automatically identify tdTomato expressing neurons.^52,53^ We used DeepLabCut (version 2.0) for automatic behavioral tracking.^40^

We performed downstream analyses using custom-written code in MATLAB and Python, using several add-on Python libraries for statistical modeling, machine learning, and data visualization. For behavioral state classification, we implemented support vector machine (SVM) decoders using the **cuml.svm.SVC** module from the RAPIDS AI framework, which provides GPU-accelerated machine learning capabilities. We evaluated classification performance using **scikit-learn’s** accuracy_score function. For regression-based analyses, we used **cuml.linear_model.Ridge** for ridge regression, enabling efficient regularized regression modeling. Data preprocessing included z-score normalization, implemented using **StandardScaler** from **scikit-learn**. To manage large datasets, we used **NumPy** for array operations and **pandas** for structured data manipulation. We visualized results using MATLAB, **matplotlib** and **seaborn**, generating heatmaps, scatter plots, and violin plots to characterize classification performances, regression results, and neural activity distributions. To apply Benjamini-Hochberg corrections for multiple comparisons, we used the MATLAB script **fdr_bh** (https://www.mathworks.com/matlabcentral/fileexchange/27418-fdr_bh*, MATLAB File Exchange*). To create violin plots, we used the code: *Bechtold, Bastian, 2016. Violin Plots for Matlab, Github Project.* https://github.com/bastibe/Violinplot-Matlab*, DOI: 10.5281/zenodo.4559847*.

#### Relevant conflicts of interest/financial disclosures

Nothing to report.

#### Funding Statement

This study was funded by the American Academy of Neurology (Neuroscience Research Training Scholarship to G.C.), Chan Zuckerberg Biohub (Physician Scientist Fellowship to G.C.), American Parkinson Disease Association (Dr. George C. Cotzias Memorial Fellowship to G.C.), Stanford Knight Initiative for Brain Resilience, Ludwig Family Foundation, and Department of Defense (Vannevar Bush Faculty Fellowship to M.J.S.).

## Notes

### Competing Interest Statement

The authors have declared no competing interest.

### Summary of Updates

The title, abstract, introduction, discussion, and overall framing have been updated to focus more specifically on amantadine and action-specific striatal neural ensembles in hypokinetic and hyperkinetic conditions. The manuscript now includes additional analyses and updated figures.

